# Optimisation of explant-specific isolation, culture, and micro-callus induction of sesame (*Sesamum indicum* L.) protoplasts

**DOI:** 10.1101/2024.10.12.618049

**Authors:** Anirban Jyoti Debnath, Debabrata Basu, Samir Ranjan Sikdar

## Abstract

The low yield of sesame (*Sesamum indicum* L.) compared to other oilseed crops hinders its successful commercialisation process. Sesame yield improvement using molecular-biotechnological tools involving gene manipulation is a better alternative than traditional breeding because it requires less time, effort, and labour. Protoplasts are plant cells devoid of the cell wall. Protoplast systems are deployed as versatile cell-based tools for tissue culture, genomics, transcriptomics, proteomics, metabolomics, and epigenetics studies leading to crop improvement. Inadequate reports of sesame protoplasts restrict its potential use in this crop. In the present study, we are reporting the successful isolation and purification of sesame protoplasts from four different sesame explants: hypocotyl, internode, leaf, and hypocotyl-derived callus using the one-step enzymatic digestion method. We have carefully optimised enzyme combinations, digestion durations, temperatures, and shaking speeds for every tested explant to obtain the highest protoplast yield and viability. Maximum yield (9.9 × 10^6^ protoplasts/gm fresh weight) and viability (92.1%) were achieved from callus explants. We purified isolated protoplasts by floating them over a 20% (w/v) sucrose solution. Among tested media, callus-derived protoplasts divided rapidly in Murashige and Skoog broth medium supplemented with 17.77 μM 6-benzylaminopurine (BAP) and 0.54 μM α-naphthalene acetic acid (NAA). In the same medium, protoplasts divided into the 2-cell stage after 2-3 days of culture and progressed to the micro-callus stage at 0.24 ± 0.05% frequency within 16-20 days of culture. Sesame protoplasts could be an excellent system for investigating potential avenues for biotechnological improvements of sesame including transient gene expression and CRISPR-based studies.

## Introduction

A protoplast is an isolated and living plant cell surrounded by the cell membrane and devoid of the cell wall. Their totipotent nature makes them a potent biotechnological tool [1]. The ability of a single protoplast to regenerate into an entire plant makes them a promising candidate for creating chimera-free plants using transgenesis studies [2]. Protoplasts are frequently used as a transient expression system in gene manipulation studies [3, 4]. Regeneration of somatic hybrids of protoplasts derived from two sexually incompatible plants can produce an entirely new plant species [5, 6]. Protoplasts serve as an excellent model system for studying various cellular processes including protein subcellular localisation, activity, transport, secretion, and interaction [7, 8], impact of (a)biotic stresses [9], growth and signal transduction [10, 11], gene expression [12, 13], and cellular and plant development processes [14, 15].

One of the oldest oilseed crops sesame (*Sesamum indicum* L., Pedaliaceae) is an excellent source of highly nutritious edible oil and seed. Voluminous presence of essential minerals like iron, magnesium, manganese, copper, and calcium, along with vitamins B1 (thiamine), E (tocopherol), and K, antioxidants like sesamol, sesamin, and sesamolin, and high amounts of unsaturated fatty acids like oleic and linoleic acids, with low amounts of saturated fatty acids such as palmitic and stearic acids make it a desirable source of nutrition [16, 17]. However, the yield of cultivated sesame is lower than other oilseeds due to undesirable traits such as inconsistent growth pattern, asynchronous capsule maturation, early pod shattering, premature senescence, and susceptibility to various biotic and abiotic stresses [16]. Addressing these issues is crucial to improve sesame production and successful commercialisation.

High-yielding protoplast isolation protocols have been successfully developed for various model plants such as tobacco (*Nicotiana tabacum* L.) [18], *Arabidopsis thaliana* L. [19] and rice (*Oryza sativa* L.) [20]. Protoplast systems have also been used as an effective tool for genetic manipulation-driven improvement in multiple plant species, including lotus, rice, clover, wheat, corn, and alfalfa [21–26]. However, reports on sesame protoplast isolation have been limited. To our knowledge, only three research groups have published their findings on sesame protoplast isolation and purification so far [27–29].

Published reports on sesame protoplasts lack sufficient detail on the protoplast isolation and purification steps. Furthermore, the success and yield of isolated protoplasts, and subsequent regeneration are highly tissue-type dependent [2]. These issues have created a challenge in sesame protoplast isolation and regeneration that has remained unresolved for 34 years since the publication of Dhingra and Batra [29]. This caveat necessitated the need to standardise a tissue-independent, detailed, and replicable sesame protoplast isolation and regeneration protocol that could be utilised to improve this ancient crop in future research.

Our article presents the first successful isolation of protoplasts from multiple sesame explants such as hypocotyl, internode, leaf, and callus. We standardised each step of protoplast isolation and purification from multiple sesame explants and induced micro-callus from hypocotyl-derived callus explants.

## Materials and methods

### Preparation of culture media, protoplast isolation enzymes, stock solutions of plant growth regulators (PGRs) and 4-morpholineethanesulfonic acid (MES)

Sesame seeds were germinated in half-strength basal MS medium (½ MS medium) [30] supplemented with 1.5% (w/v) sucrose and 0.6% (w/v) agar. Full-strength basal MS medium supplemented with 3% sucrose (w/v) and 0.8% agar (w/v) was used for clonal propagation and callus culture. The divisional response of protoplasts was studied using basal MS broth medium supplemented with 5 mM of MES and 0.6 M glucose. MS medium, sucrose and agar were purchased from HiMedia (Mumbai, India). MES was purchased from Sigma-Aldrich (USA). Sterilisation of media was carried out by autoclaving at 121°C temperature with 15 psi pressure for 15 minutes, except protoplast culture media, which were filter sterilised by passing through a filter paper having a porosity of 0.22 µm. For protoplast isolation, cellulase “Onozuka” R-10, cellulase “Onozuka” RS, macerozyme R-10 (Yakult Pharmaceutical Industry Co. Ltd., Japan), sumizyme 6,000, sumizyme 54,000 (Shin Nihon Chemicals Co. Ltd., Japan), driselase, and lysing enzyme (Sigma-Aldrich, USA) were used. Enzyme solutions were prepared in 0.6 M mannitol (HiMedia, Mumbai, India) supplemented with 0.2% (w/v) CaCl_2_.2H_2_O (HiMedia, Mumbai, India) and filter sterilised. Adjustment of pH in all the media and enzyme solutions was made before autoclaving or filter sterilisation at 5.8 by the suitable addition of 1N HCl and 1N KOH solutions. BAP, NAA, and 2,4-D stock solutions were prepared by using 1N NaOH as a solvent. The concentration of all the stock solutions of PGRs (Sigma-Aldrich, USA) was maintained at 1 mg/ml. MES was dissolved in double distilled water to make a 1 M stock solution. All the prepared stock solutions were filter sterilised and stored at 4°C in an amber bottle. PGRs were added to the media before autoclaving.

### Plant material, seed culture, clonal propagation, and callus culture

*Sesamum indicum* L. cultivar JK-1 seeds were collected from the National Bureau of Plant Genetic Resources, Pusa, New Delhi, India. Healthy and uniform seeds were washed with 50% commercial bleach (RIN ALA, Hindustan Unilever Limited, India) containing 4% chlorine (v/v) for 5 min. Surface sterilisation of the cleaned seeds was done with 0.1% (w/v) aqueous mercuric chloride solution for 2.5 min followed by 70% ethanol for 10 s. Subsequently, surface-sterilised seeds were washed four times with sterile distilled water, transferred to Petri dishes having 25 ml sterilised ½ MS medium, and incubated at 28±2°C in the dark. Five weeks old hypocotyls were used for protoplast isolation (Fig. 1a).

**Fig. 1.**
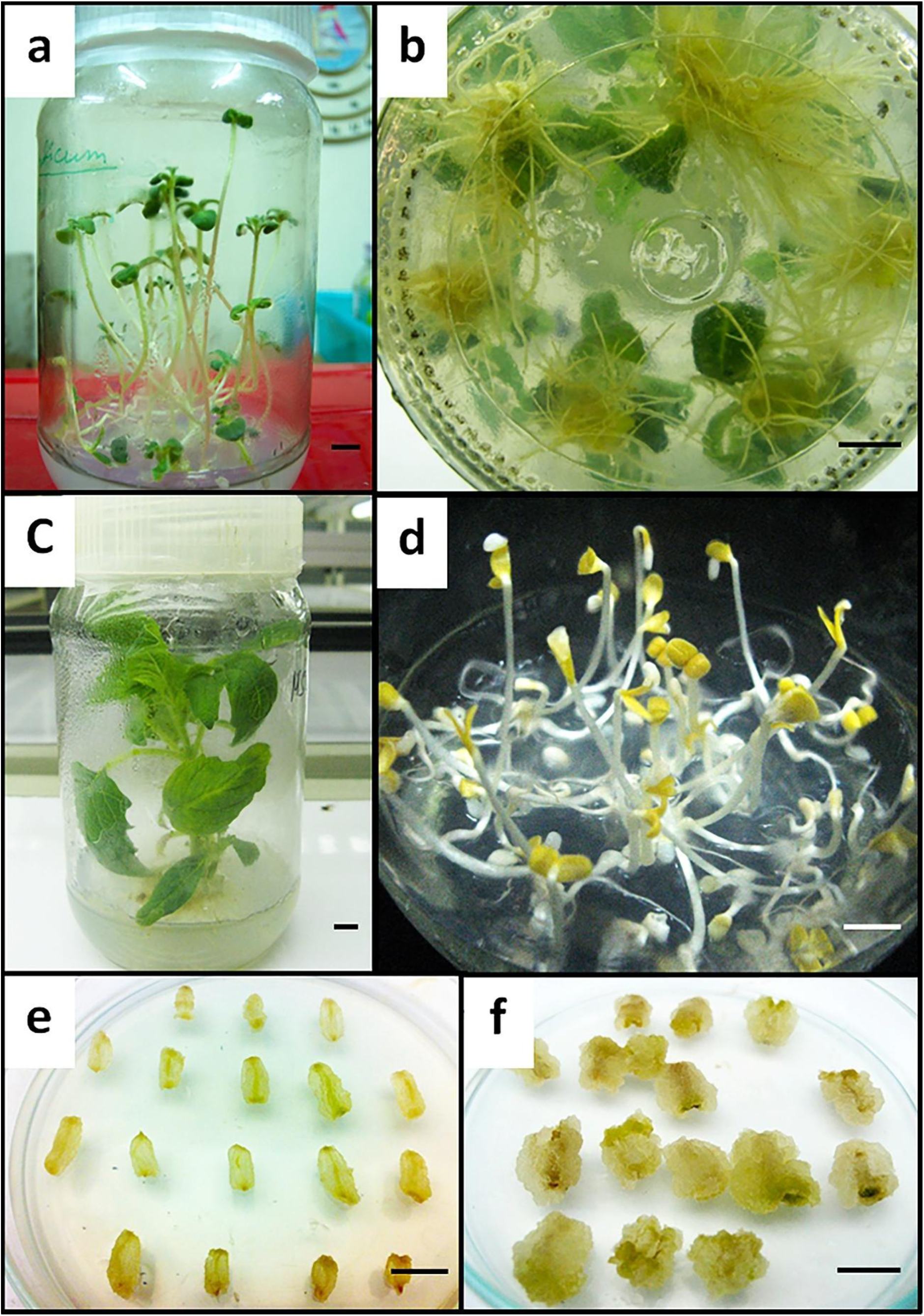
Sesame explants used for protoplast isolation; sesame seeds were germinated in half-strength basal MS medium supplemented with 1.5% (w/v) sucrose and 0.6% (w/v) agar **a** hypocotyl explants from 5-week-old seedlings were used for protoplast isolation; **b** roots were induced from apical shoot tip explants in basal MS medium fortified with 2.69 µM of NAA to initiate clonal propagation; **c** a five-week-old clonally propagated culture from which leaf and internode explants were used for protoplast isolation; **d** hypocotyls from 15-day-old seedlings were used to initiate callus culture on basal MS medium supplemented with 1.13 µM 2,4-D, 5.37 µM NAA, and 3.33 µM BAP (MS-5) medium; **e and f** two-stage callus induction from hypocotyl explants for protoplast isolation; **e** in stage-I, 15-day-old hypocotyl explants were cultured in MS-5 medium for 5 days; **f** calluses from MS-5 medium were inoculated on basal MS medium supplemented with 0.54 µM NAA, and 17.74 µM BAP (S-5) medium, cultured for another 15 days to induce stage-II calluses, and used as explants for protoplast isolation; bars 10 mm

The clonally propagating population of sesame was established by root induction of apical shoot tips from germinated seedlings in MS medium supplemented with 2.69 µM of NAA (Fig. 1b), and subsequent transfer of rooted explants in fresh basal MS medium. Internodes and leaves from four to five-week-old clonally propagated cultures were used for protoplast isolation (Fig. 1c).

We initiated stage-I of the two-stage callus induction by culturing 15-day-old hypocotyls (Fig. 1d) in basal MS medium supplemented with 1.13 µM 2,4-D, 5.37 µM NAA, and 3.33 µM BAP (MS-5) (Fig. 1e). After five days of culture, these calluses were inoculated in basal MS medium supplemented with 0.54 µM NAA, and 17.74 µM BAP (S-5) to induce the stage-II calluses (Fig. 1f). We designated stage-II calluses simply as “callus”. Stage-II calluses were used for protoplast isolation after 15 days of culture in the S-5 medium. All the clonally propagated and callus cultures were incubated in the culture room having a temperature of 25±2°C, and relative humidity of 60-65% under 16/8 photoperiod having an irradiance of 35 µmole/m^2^/s photosynthetic photon flux density using daylight fluorescent tube lights (Philips Champion, India).

### Isolation of protoplasts

Protoplasts were isolated by one step [31] enzymatic method [32] from the hypocotyl, internode, leaf, and hypocotyl-derived callus of sesame. About one gm sesame explant tissue was finely chopped with a sharp scalpel blade, placed in the ten ml enzyme solution, and incubated for protoplast isolation. Enzyme combination and incubation parameters were standardised for each explant in the following three experiments. We adopted two leaf-derived protoplast isolation protocols depending on the incubation time – leaf “fast” and leaf “slow”. Individual protocols were designed for the rest of the explants.

In the first experiment, we studied the combinational effect of enzyme incubation period temperature and duration on protoplast isolation. We applied different enzyme combinations for each explant type (Supplementary Table S1). Shaking was not applied. The combined treatments of incubation period temperature and duration ranged from 16–36°C and 1–18 hrs., respectively. The treatments for hypocotyl, internode, leaf “fast”, leaf “slow”, and callus explants were designated as HTT1–HTT9, ITT1–ITT9, LFTT1–LFTT9, LSTT1– LSTT9, and CTT1–CTT9, respectively.

In the second experiment, the effect of shaking on protoplast isolation was observed. We applied 0, 60 and 100 rpm shaking treatment using a rotary shaker during incubation. The rest of the protoplast isolation parameters were kept unchanged as previously. The treatments for hypocotyl, internode, leaf “fast”, leaf “slow”, and callus explants were designated as HS1–HS3, IS1–IS3, LFS1–LFS3, LSS1–LSS3, and CS1–CS3, respectively.

In the third experiment, we studied the effect of enzyme combinations on protoplast isolation. We used various combinations of 0.5–3.5% cellulase R-10, 0.25–0.5% cellulase RS, 1.0–2.0% macerozyme R-10, 0.25–2.0% sumizyme 54,000, 0.25–2.0% sumizyme 6,000, 0.25–0.5% driselase, and 0.25–1.0% lysing enzyme in this experiment. The rest of the protoplast isolation parameters were kept unchanged as previously. The treatments for hypocotyl, internode, leaf “fast”, leaf “slow”, and callus explants were designated as HE1– HE9, IE1–IE9, LFE1–LFE9, LSE1–LSE9, and CE1–CE9, respectively.

### Purification of protoplasts

Following the incubation period, the digested tissue containing protoplasts was passed through a steel sieve having a pore size of 100 µM to isolate the debris. The sieved enzyme-protoplast solution was centrifuged at 100 x *g* for three minutes, and the supernatant was decanted. The pellet containing protoplast was dissolved in the washing solution (0.6 M mannitol supplemented with 0.2% (w/v) CaCl_2_.2H_2_O, pH 5.8) and gradually layered on the top of 9 ml of 20% (w/v) sucrose solution taken in a 15 x 125 mm screw-cap centrifuge tube. It was then centrifuged at 100 x *g* for ten minutes. The intact protoplasts formed a prominent ring on the upper phase of the sucrose solution. The ring was collected using a glass Pasteur pipette. The collected purified protoplast solution was then re-suspended in fresh washing solution in a 9:1 ratio and was washed by centrifugation at 100 x g thrice. Finally, the pellet of purified and washed protoplasts was re-suspended in 1 ml of washing solution. Yield was counted using a Haemocytometer and an inverted microscope (Fluovert, Leitz, Germany), and represented as the number of protoplasts/gm of fresh weight (FW) of the explant tissue; the solution was used for downstream experiments.

### Checking for the viability of the purified protoplasts

The viability of the purified protoplasts was tested using the fluorescein diacetate (FDA) (Sigma-Aldrich, USA) staining method [116] using the following protocol. The 0.5% (w/v) stock of FDA staining solution was prepared in acetone and diluted to 0.01% with washing solution. The diluted stock of staining solution was mixed with an equal volume of freshly isolated protoplast solution and incubated for five minutes in the dark. The protoplasts were observed under a fluorescent microscope (Fluovert, Leitz, Germany) using an excitation wavelength of 365 nm. The number of viable yellow-green fluorescent and non-viable, non-fluorescent protoplasts were counted using a Haemocytometer. Protoplast viability was calculated by the following formula:

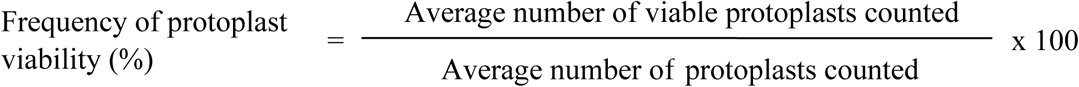

### Culture of purified protoplasts

Protoplasts were cultured in basal MS broth medium fortified with all possible combinations of 0.54 µM or 1.08 µM of NAA, and 4.44 µM, 11.11 µM, 17.77 µM or 33.33 µM of BAP (medium 1–8). For each culture, 2 ml of medium was used. The plating density was 5 × 10^4^ cells/ml for all the cultures. Culture plates were incubated at 28 ± 2°C in the dark. All cultures were refreshed with the same media once a week.

### Data recording method and statistical analyses

The protoplast yield and viability data were averaged from three sets of experiments. Divisional response data was recorded after three weeks of protoplast culture. The appearance of each of the 2-celled, 4-celled, 8-celled, micro-colony, and micro-callus stages was counted for all the treatments independently in several microscopic fields. The average data of 10 such fields for each divisional stage in the treatments represented the final data of an experiment. Three such experiments were performed to carry out further statistical analysis. All the data are represented as mean ± standard error (SE). Data were analysed by one-way ANOVA (Analysis of Variance) to calculate statistical significance. The significant difference in treatment data was assessed through Duncan’s multiple range test at *P* < 0.05 [33]. All the analyses were performed in SPSS version 19 (IBM Corporation, USA).

## Results

We isolated protoplasts from hypocotyl, internode, leaf, and hypocotyl-derived callus explants of sesame. Callus explants were prepared in two stages. Hypocotyl-derived MS-5 medium-cultured stage-I calluses were subsequently inoculated in S-5 medium to induce the stage-II calluses. The calluses in both media were swollen and fluffy with 100% induction frequency (Fig. 1e, f). The stage-II calluses were used for protoplast isolation.

### Effect of enzyme incubation period temperature and duration on protoplast isolation

We used different pre-determined enzyme combinations for each tested explant type, and static shaking during incubation in this experiment. Results depicted significant variations in optimum incubation temperature and duration explant-wise (Table 1). From the hypocotyl explants, protoplast yield (3.54 ± 0.01 × 10^6^ protoplasts/gm FW) and viability (51.99 ± 1.57%) were highest at 25°C and 16 hrs. incubation (HTT5). Using internode explants, we achieved maximum protoplast yield (3.46 ± 0.26 × 10^6^ protoplasts/gm FW) and viability (50.72 ± 1.39%) at 27°C and 16 hrs. incubation (ITT8). The mesophyll protoplast release was quick in the “fast” protocol; leaf tissues were incubated at 32°C for 2.5 hrs. (LFTT5) to achieve an optimal yield (4.38 ± 0.25 × 10^6^ protoplasts/gm FW) and viability (56.93 ± 1.02%). Interestingly, in the leaf “slow” protocol, both yield (6.77 ± 0.20 × 10^6^ protoplasts/gm FW) and viability (80.16 ± 0.73%) of mesophyll protoplasts increased dramatically with a reduced incubation temperature of 16°C and 16 hrs. incubation period (LSTT2). Among all the tested explants, we obtained the highest yield from callus explants (8.13 ± 0.33 × 10^6^ protoplasts/gm FW) and viability (80.43 ± 0.68%) by incubating them at 22°C for 16 hrs. (CTT2).

**Table 1.**
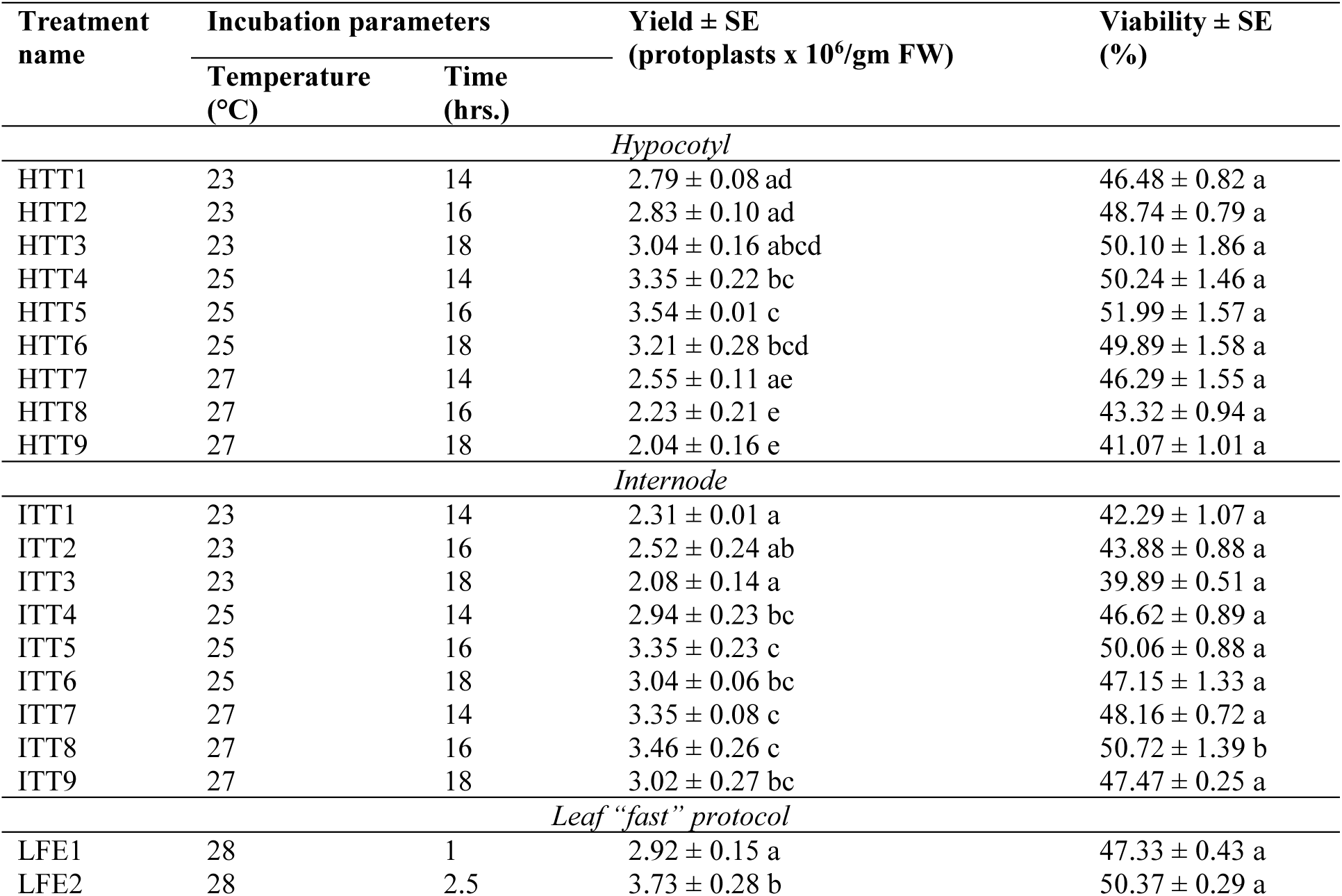
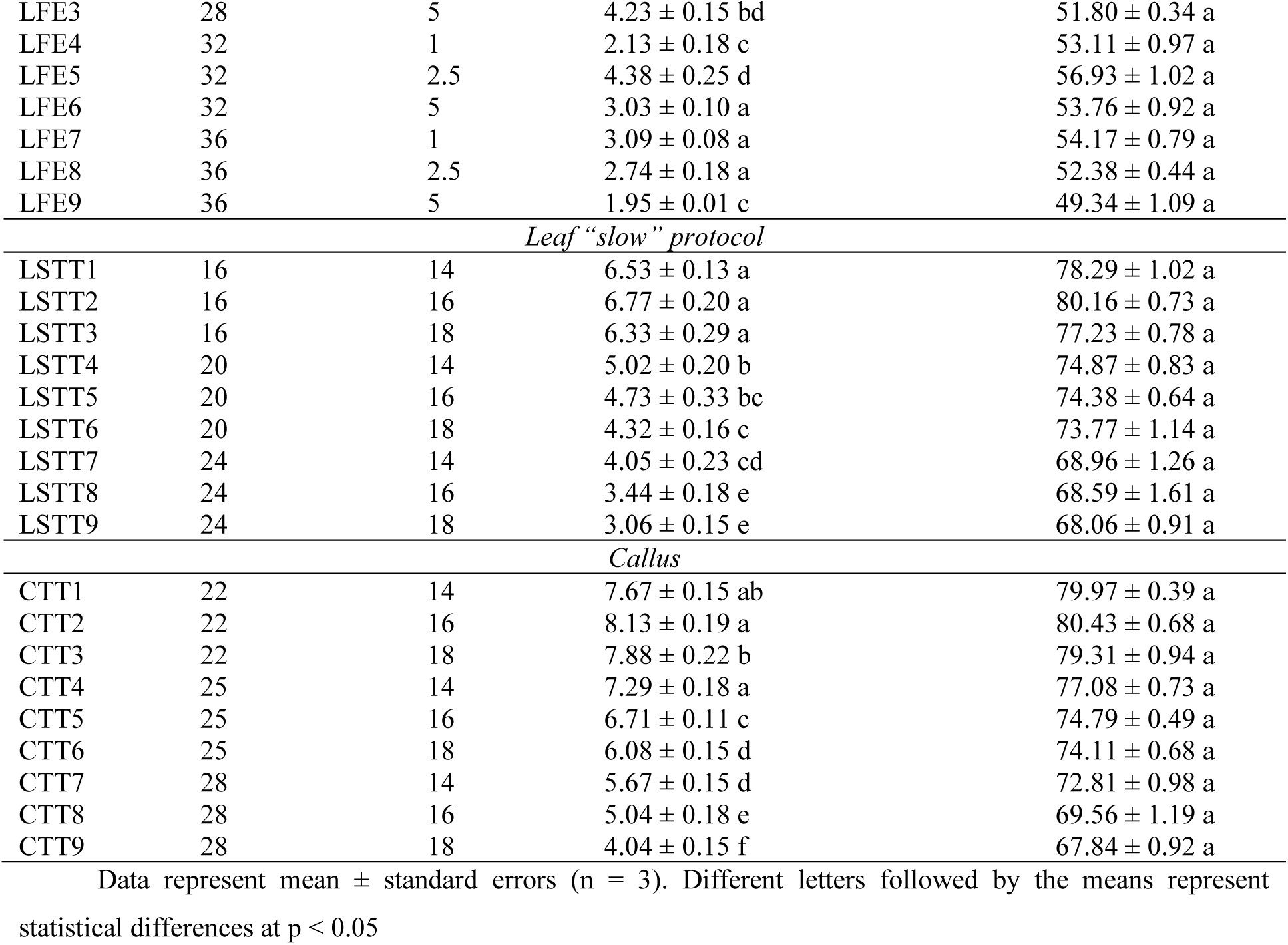
Effect of various combinations of incubation temperature and time on the yield and viability of sesame protoplast isolation from different explants.

### Effect of shaking during enzyme incubation on protoplast isolation

In this experiment, we used unaltered enzyme combinations and optimised incubation temperature and duration found in the previous experiment. We used 0, 60, and 100 rpm incubation shaking for this analysis. Shaking did not influence protoplast isolation in all explants (Table 2). The 60 rpm (HS2) and 100 rpm (LFS3) shaking improved protoplast isolation efficiency only from hypocotyl (yield: 3.79 ± 0.25 × 10^6^ protoplasts/gm FW; viability: 53.12 ± 1.26%) and leaf explants with “fast” protocol (yield: 6.04 ± 0.23 × 10^6^ protoplasts/gm FW; viability: 67.17 ± 0.84%), respectively. Shaking did not benefit other treatments. Among all the tested explants, callus-derived protoplast yield (8.13 ± 0.33 × 10^6^ protoplasts/gm FW) and viability (80.43 ± 0.68%) using 0 rpm shaking (CS1) was highest.

**Table 2.** Effect of shaking during the incubation period on the yield and viability of sesame protoplast isolation from different explants.

| Treatment name | Shaking (RPM) | Yield ± SE (protoplasts x 10<sup>6</sup>/gm FW) | Viability ± SE (%) |
| --- | --- | --- | --- |
| *Hypocotyl* |  |  |  |
| HS1 | 0 | 3.59 ± 0.09 ab | 50.28 ± 0.68 a |
| HS2 | 60 | 3.79 ± 0.14 b | 53.12 ± 1.26 a |
| HS3 | 100 | 3.11 ± 0.18 a | 45.30 ± 0.78 a |

| *Internode* |  |  |  |
| --- | --- | --- | --- |
| IS1 | 0 | 5.81 ± 0.44 a | 50.01 ± 0.92 a |
| IS2 | 60 | 5.04 ± 0.44 a | 43.61 ± 1.36 a |
| IS3 | 100 | 4.85 ± 0.44 a | 39.97 ± 1.13 a |
| *Leaf “fast” protocol* |  |  |  |
| LFS1 | 0 | 4.32 ± 0.15 a | 56.78 ± 1.77 a |
| LFS2 | 60 | 5.45 ± 0.20 b | 65.09 ± 0.66 a |
| LFS3 | 100 | 6.04 ± 0.13 c | 67.17 ± 0.84 a |
| *Leaf “slow” protocol* |  |  |  |
| LSS1 | 0 | 6.74 ± 0.18 a | 80.07 ± 0.81 a |
| LSS2 | 60 | 5.28 ± 0.24 b | 69.19 ± 1.17 a |
| LSS3 | 100 | 4.93 ± 0.26 b | 64.47 ± 0.74 a |
| *Callus* |  |  |  |
| CS1 | 0 | 8.17 ± 0.25 a | 80.88 ± 1.04 a |
| CS2 | 60 | 7.08 ± 0.18 b | 73.08 ± 0.93 a |
| CS3 | 100 | 6.54 ± 0.27 b | 70.91 ± 1.28 a |

Data represent mean ± standard errors (n = 3). Different letters followed by the means represent statistical differences at $p < 0.05$

### Effect of enzyme combination on protoplast isolation

We used different enzyme combinations to find the tested explant-wise optima of protoplast isolation efficiency (Table 3). We applied optimised enzyme incubation temperature, duration, and shaking speed for each explant type found in previous experiments. The enzyme solution containing 0.5% cellulase R-10, 0.5% cellulase RS, 0.25% driselase, and 1.0% macerozyme R-10 (HE2) provided the highest yield (4.75 ± 0.08 × 10^6^ protoplasts/gm FW) and viability (67.15 ± 0.42%) from the hypocotyl explants. For the internode explants, the combination of 3.5% cellulase R-10, 0.5% cellulase RS, and 1.0% macerozyme R-10 (IE4) maximised the yield (5.44 ± 0.20 × 10^6^ protoplasts/gm FW) and viability (68.37 ± 0.28%). Sumizymes and the lysing enzyme were crucial for our sesame mesophyll protoplast isolation because the yield was negligible without them. For the “fast” protocol of mesophyll protoplast release, a mixture of 0.25% cellulase R-10, 2.0% sumizyme 54,000, 2.0% sumizyme 6,000, 0.25% lysing enzyme, and 1.0% macerozyme R-10 (LFE4) provided maximum yield (6.91 ± 0.15 × 10^6^ protoplasts/gm FW) and viability (82.16 ± 1.18%). Milder enzymatic treatment was needed for the “slow” protocol for mesophyll protoplast isolation. In this method, a solution of 1.00% cellulase R-10, 0.5% sumizyme 54,000, 0.5% sumizyme 6,000, and 1.0% macerozyme R-10 (LSE3) provided the optimal yield (7.15 ± 0.13 × 10^6^ protoplasts/gm FW) and viability (86.76 ± 0.78%). We achieved the highest yield (9.88 ± 0.19 × 10^6^ protoplasts/gm FW) and viability (92.05 ± 0.81%) of sesame protoplasts from callus explants using 1.5% cellulase R-10, 0.5% driselase, and 1.0% macerozyme R-10 (CE3).

**Table 3.** Effect of enzyme combinations during the incubation period on the yield and viability of sesame protoplast isolation from different explants.

| Treatment name | Enzyme combinations | Yield ± SE<br>(protoplasts $\times 10^6$/gm FW) | Viability ± SE<br>(%) |
| --- | --- | --- | --- |
| *Hypocotyl* |  |  |  |
| HE1 | Cellulase R-10 0.5%<br>Cellulase RS 0.25%<br>Driselase 0.25%<br>Macerozyme R-10 1.0% | 3.71 ± 0.12 a | 53.46 ± 0.25 a |
| HE2 | Cellulase R-10 0.5%<br>Cellulase RS 0.5%<br>Driselase 0.25%<br>Macerozyme R-10 1.0% | 4.75 ± 0.08 b | 67.15 ± 0.42 b |
| HE3 | Cellulase R-10 1.0%<br>Cellulase RS 0.25%<br>Driselase 0.5%<br>Macerozyme R-10 2.0% | 3.03 ± 0.22 c | 45.53 ± 0.31 c |
| HE4 | Cellulase R-10 1.0%<br>Cellulase RS 0.5%<br>Driselase 0.5%<br>Macerozyme R-10 2.0% | 2.61 ± 0.28 c | 41.02 ± 0.24 c |
| *Internode* |  |  |  |
| IE1 | Cellulase R-10 0.5%<br>Cellulase RS 0.25%<br>Driselase 0.25%<br>Macerozyme R-10 1.0% | 3.42 ± 0.21 a | 49.12 ± 1.12 a |
| IE2 | Cellulase R-10 2.0%<br>Cellulase RS 0.5%<br>Driselase 0.5%<br>Macerozyme R-10 2.0% | 4.79 ± 0.27 b | 54.63 ± 0.95 ab |
| IE3 | Cellulase R-10 2.0%<br>Cellulase RS 0.5%<br>Driselase 0.0%<br>Macerozyme R-10 2.0% | 5.23 ± 0.18 b | 61.13 ± 0.78 ab |
| IE4 | Cellulase R-10 3.5%<br>Cellulase RS 0.5%<br>Driselase 0.0%<br>Macerozyme R-10 1.0% | 5.44 ± 0.20 b | 68.37 ± 0.28 b |
| *Leaf "fast" protocol* |  |  |  |
| LFE1 | Cellulase R-10 1.0%<br>Sumizyme 54,000 1.0%<br>Sumizyme 6,000 1.0%<br>Lysing enzyme 1.0%<br>Macerozyme R-10 1.0% | 5.95 ± 0.23 a | 66.12 ± 1.29 a |
| LFE2 | Cellulase R-10 2.0%<br>Sumizyme 54,000 2.0%<br>Sumizyme 6,000 2.0%<br>Lysing enzyme 1.0%<br>Macerozyme R-10 2.0% | 6.71 ± 0.21 b | 77.07 ± 0.64 a |
| LFE3 | Cellulase R-10 0.25%<br>Sumizyme 54,000 2.0%<br>Sumizyme 6,000 2.0%<br>Lysing enzyme 1.0%<br>Macerozyme R-10 1.0% | $6.33 \pm 0.19$ ab | $72.13 \pm 0.68$ a |
| LFE4 | Cellulase R-10 0.25%<br>Sumizyme 54,000 2.0%<br>Sumizyme 6,000 2.0%<br>Lysing enzyme 0.25%<br>Macerozyme R-10 1.0% | $6.91 \pm 0.15$ b | $82.16 \pm 1.18$ a |
| *Leaf "slow" protocol* |  |  |  |
| LSE1 | Cellulase R-10 1.0%<br>Sumizyme 54,000 1.0%<br>Sumizyme 6,000 0.5%<br>Lysing enzyme 0.25%<br>Macerozyme R-10 1.0% | $6.71 \pm 0.25$ a | $79.33 \pm 0.47$ a |
| LSE2 | Cellulase R-10 0.25%<br>Sumizyme 54,000 0.5%<br>Sumizyme 6,000 0.5%<br>Lysing enzyme 0.25%<br>Macerozyme R-10 2.0% | $5.45 \pm 0.12$ b | $57.43 \pm 1.02$ b |
| LSE3 | Cellulase R-10 1.0%<br>Sumizyme 54,000 0.5%<br>Sumizyme 6,000 0.5%<br>Lysing enzyme 0.0%<br>Macerozyme R-10 1.0% | $7.15 \pm 0.13$ a | $86.76 \pm 0.78$ a |
| LSE4 | Cellulase R-10 2.0%<br>Sumizyme 54,000 0.25%<br>Sumizyme 6,000 0.25%<br>Lysing enzyme 0.25%<br>Macerozyme R-10 1.0% | $5.72 \pm 0.24$ b | $61.43 \pm 0.63$ b |
| *Callus* |  |  |  |
| CE1 | Cellulase R-10 0.5%<br>Driselase 0.5%<br>Macerozyme R-10 1.0% | $8.04 \pm 0.22$ a | $80.25 \pm 1.46$ a |
| CE2 | Cellulase R-10 1.0%<br>Driselase 0.25%<br>Macerozyme R-10 2.0% | $8.63 \pm 0.07$ ab | $76.41 \pm 1.09$ a |
| CE3 | Cellulase R-10 1.5%<br>Driselase 0.5%<br>Macerozyme R-10 1.0% | $9.88 \pm 0.19$ c | $92.05 \pm 0.81$ a |
| CE4 | Cellulase R-10 2.0%<br>Driselase 0.25%<br>Macerozyme R-10 2.0% | $9.08 \pm 0.23$ b | $88.51 \pm 0.58$ a |

Data represent mean ± standard errors (n = 3). Different letters followed by the means represent statistical differences at $p < 0.05$

### Optimised conditions for protoplast isolation and purification from different explants

The optimised protocols for protoplast isolation described above are summarised as follows for different tested explants.

### Hypocotyls

The finely chopped hypocotyl explants from 5-week-old seedlings (Fig. 1a) were placed in the enzyme mixture of 0.5% cellulase R-10, 0.5% cellulase RS, 0.25% driselase, and 1.0% macerozyme R-10. The mixture was incubated at 25°C with 60 rpm shaking for 16 hrs. Fig. 2a represents freshly isolated hypocotyl-derived protoplasts.

**Fig. 2.**
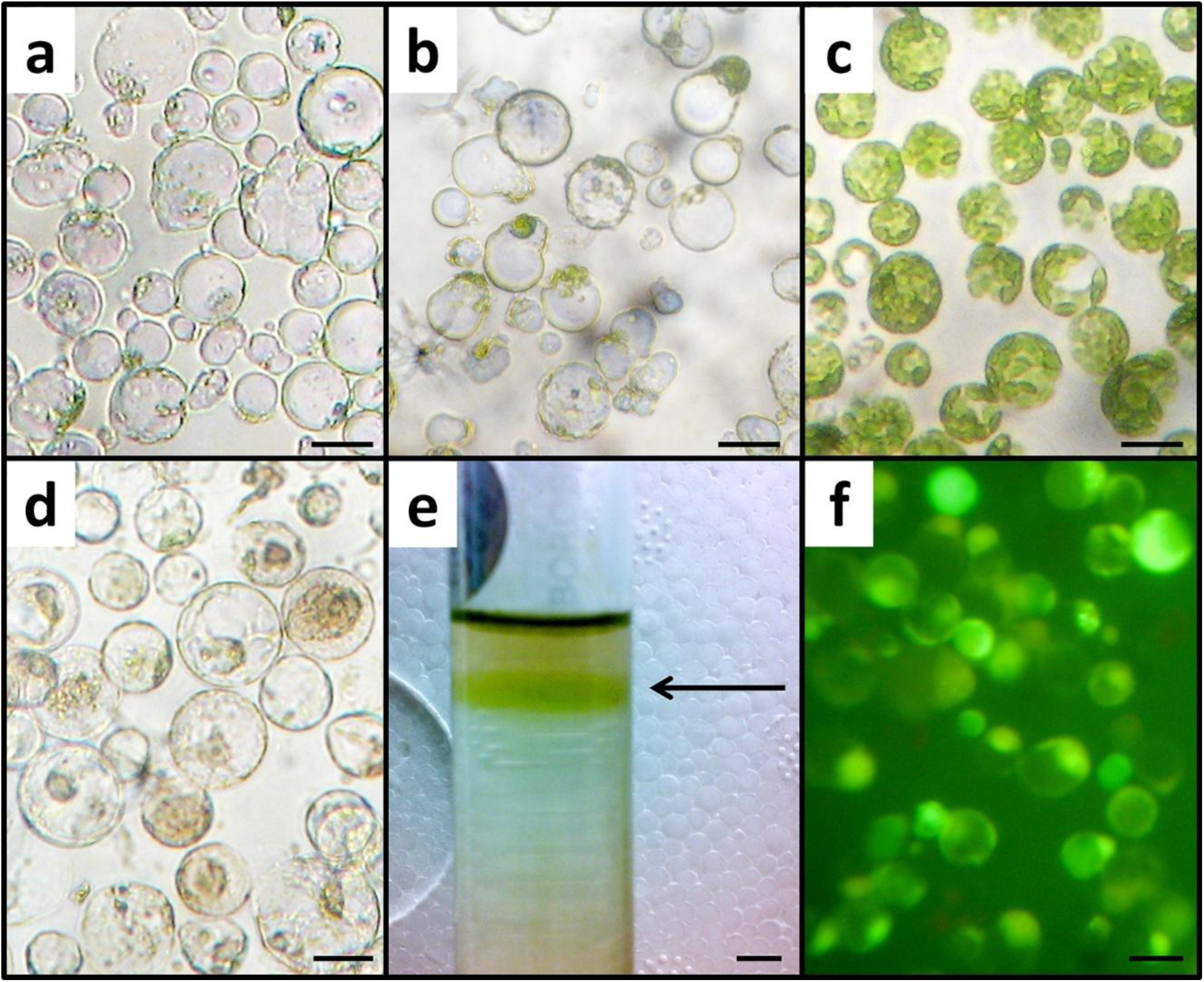
Protoplasts from different sesame explants; about one gm sesame explant tissue was finely chopped with a sharp scalpel blade, placed in the ten ml enzyme solution, and incubated for protoplast isolation; freshly isolated protoplasts from **a** hypocotyl **b** internode **c** leaf and **d** callus explants; **e** isolated healthy protoplasts formed a ring (black arrow) on the upper phase of the 20% sucrose solution during purification step; **f** isolated and purified protoplasts were stained with FDA for viability assessment; alive protoplasts fluoresce green when excited at 365 nm; bars 50 μm (**a**, **b**, **c**, **d**, and **f**) and 5 mm (**e**)

### Internodes

We used basal MS medium fortified with 2.69 µM of NAA for root induction from the apical shoot tip explants to initiate clonal propagation (Fig. 1b). Internode explants from the four to five-week-old clonally propagated culture were finely chopped and placed in the mixture of 3.5% cellulase R-10, 0.5% cellulase RS, and 1.0% macerozyme R-10 and incubate at 27°C without shaking for 16 hrs. Fig. 2b represents freshly isolated internode-derived protoplasts.

### Leaf “fast” protocol

Mature leaves from the four to five-week-old clonally propagated cultures (Fig. 1c) were chopped finely and incubated in the enzyme combination of 0.25% cellulase R-10, 2.0% sumizyme 54,000, 2.0% sumizyme 6,000, 0.25% lysing enzyme, and 1.0% macerozyme R-10 at 32°C with 100 rpm shaking for 2.5 hrs.

### Leaf “slow” protocol

Mature leaves from the four to five-week-old clonally propagated cultures (Fig. 1c) were chopped finely and incubated in the enzyme combination of 1.0% cellulase R-10, 0.5% sumizyme 54,000, 0.5% sumizyme 6,000, 1.0% macerozyme R-10 at 16°C for 16 hrs without shaking. Fig. 2c represents freshly isolated mesophyll protoplasts.

### Callus

The 15-day-old hypocotyls were cultured in the MS-5 medium for five days to induce stage-I calluses. The stage-I calluses were inoculated in the S-5 medium. After 15 days of culture in the S-5 medium, stage-II calluses were finely chopped and incubated in the mixture of 1.5% cellulase R-10, 0.5% driselase, and 1.0% macerozyme R-10 at 22°C for 16 hrs without shaking. Fig. 2d represents freshly isolated callus-derived protoplasts.

Isolated protoplasts from every explant were purified by floating over a 20% sucrose solution. Intact protoplasts formed a ring in the upper phase of the solution (Fig. 2e). Purified protoplasts were washed in the washing solution (0.6 M mannitol supplemented with 0.2% (w/v) CaCl_2_.2H_2_O, pH 5.8). FDA-stained viable protoplasts appeared green under the fluorescence microscope (Fig. 2f). The protoplast isolation and purification steps have been summarised as a flow chart in Fig. 3.

**Fig. 3.**
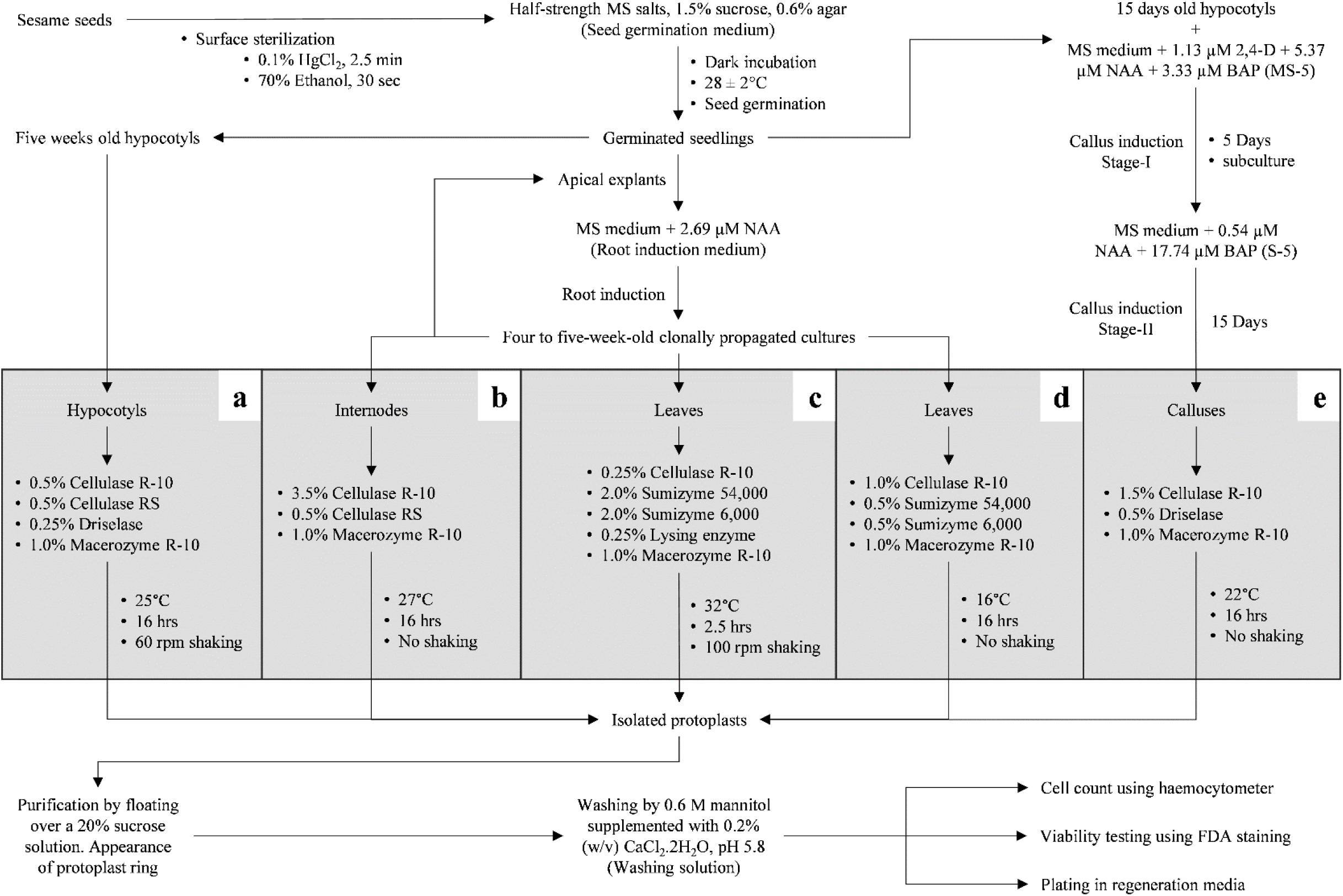
Flow diagram of protoplast isolation and purification from different sesame explants; the diagram represents optimised steps of sesame protoplast isolation and purification protocol starting from seed germination via **a** hypocotyl, **b** internode, **c** leaf “slow”, **d** leaf “fast”, and **e** callus explant preparations up to cell count, viability testing, and plating stage

### Divisional response of protoplasts

We cultured the hypocotyl, internode, and leaf-derived protoplasts in basal MS broth medium supplemented with various combinations of 2,4-D, NAA, and BAP. However, after several attempts, protoplasts obtained from these explants did not divide after the 2-cell stage (data not shown). We studied the callus-derived protoplasts for the divisional response using basal MS broth medium fortified with different combinations of NAA and BAP (medium-1 to medium-8). The callus-derived protoplasts started dividing in all treatment media (Fig. 4a, b, c). Within a couple of days of culture, a microtubule-like structural arrangement (Fig. 4a) appeared in the dividing cells followed by a septa-like structure at the divisional plane (Fig. 4b). Two-celled stage appeared within 2-3 days of culture (Fig. 4c). Among all the tested PGR combinations in the culture media, the highest frequency of 6.16 ± 0.09% of two-cell stage was found in the medium-3 (MS medium supplemented with 0.54 μM NAA and 17.77 μM BAP), followed by 5.78 ± 0.21% in the medium-4 (MS medium supplemented with 0.54 μM NAA and 33.33 μM BAP) (Table 4). The 4-cell stage appeared only in three media within 6-8 days of culture (Fig. 4d). The maximum frequency of 1.91 ± 0.17% of this stage was found in the medium-3 (MS basal medium supplemented with 0.54 μM NAA and 17.77 μM BAP), followed by 0.59 ± 0.07% in the medium-4 (MS basal medium supplemented with 0.54 μM NAA and 33.33 μM BAP), and 0.55 ± 0.06% in the medium-2 (MS basal medium supplemented with 0.54 μM NAA and 11.11 μM BAP). We found the appearance of 8-cell, 16-cell, micro-colony, and micro-callus stages only in the medium-3 (Fig. 4e, f, h, i, j). In this medium, micro-colonies appeared within 14-16 days of culture at the divisional rate of 0.47 ± 0.06% (Fig. 4g, h). Micro-calluses were observed in the same medium at the frequency of 0.24 ± 0.05% after 18-20 days of culture (Fig. 4i, j). Protoplast division seized at this stage. After several attempts, micro-colonies did not grow further.

**Fig. 4.**
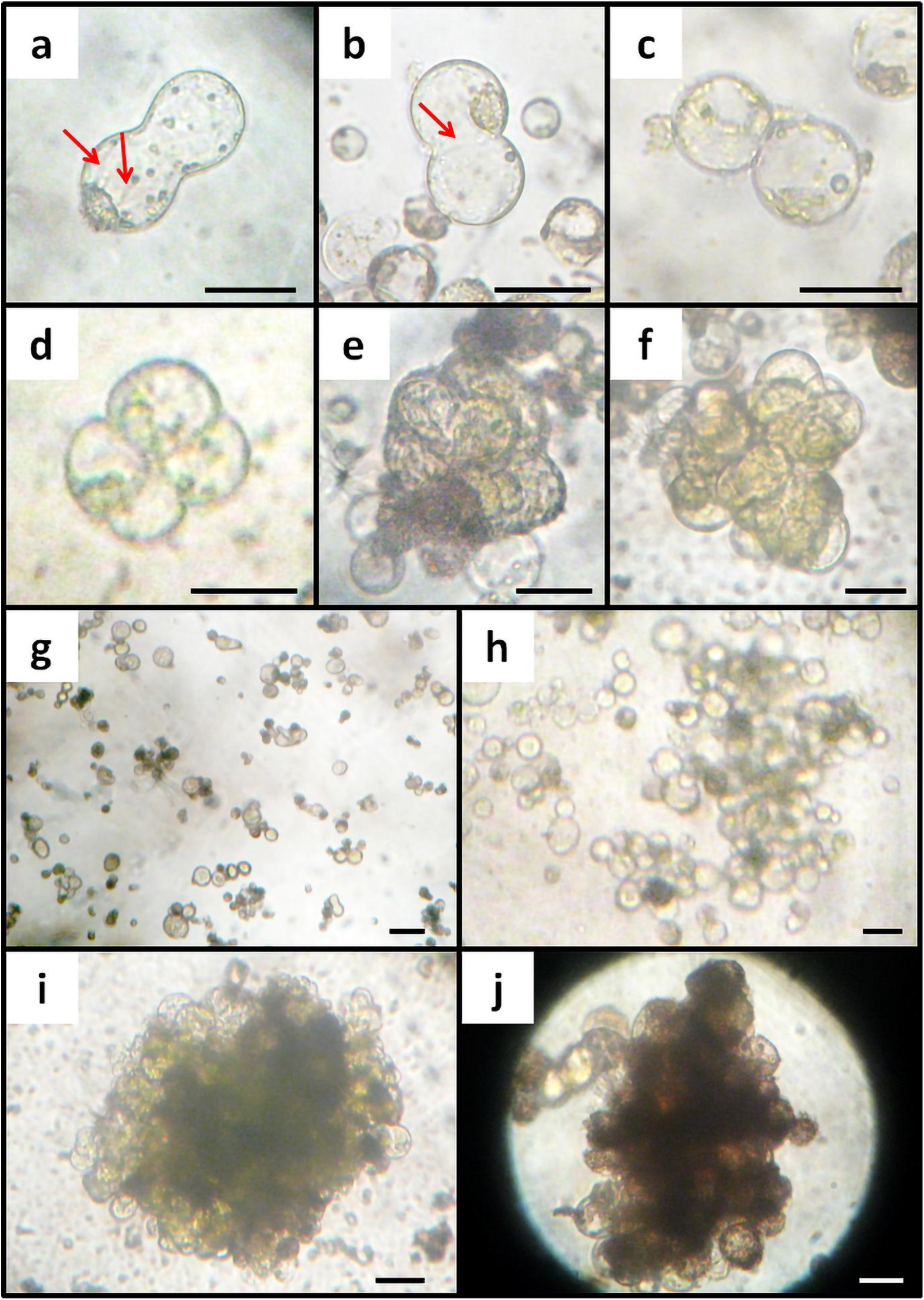
Cell division of sesame callus-derived protoplasts; optimum divisional response from sesame protoplasts was obtained when they were cultured in 2 ml basal MS broth medium supplemented with 0.54 μM NAA and 17.77 μM BAP (medium-3) at the plating density of 5 × 10^4^ cells/ml; cultures were refreshed weekly with the same medium; before the first division of protoplasts, appearance of **a** microtubule-like structural arrangement (arrow-head), and **b** a septa-like structure at the divisional plane (arrow-head) was observed within a couple of days of culture; **c** 2-cell stage appeared within 2-3 days of culture; **d** 4-cell stage appeared within 6-8 days of culture; **e** 8-cell stage, **f** 16-cell stage, and **g** micro-colonies appeared within 14-16 days of culture; **h** magnified view of a micro-colony; **i** micro-calluses appeared after 18-20 days of culture; **j** magnified view of a micro-callus; bars 50 μm (**a**, **b**, **c**, **d**, **e**, **f**, and **j**) and 100 μm (**g**, **h**, and **i**)

**Table 4.** The divisional responses of sesame callus-derived protoplasts in different protoplast culture medium.

| Protoplast Culture Medium | Amount of hormone | | Divisional frequency (%) at different divisional stage ± SE | | | | |
| --- | --- | --- | --- | --- | --- | --- | --- |
| | BAP (μM) | NAA (μM) | 2-cell stage | 4-cell stage | 8-cell stage | Micro-colony | Micro-callus |
| Medium-1 | 4.44 | 0.54 | 4.12 ± 0.18 ab | 0 ± 0 a | 0 ± 0 a | 0 ± 0 a | 0 ± 0 a |
| Medium-2 | 11.11 | 0.54 | 3.82 ± 0.17 ab | 0.55 ± 0.06 a | 0 ± 0 a | 0 ± 0 a | 0 ± 0 a |
| Medium-3 | 17.77 | 0.54 | 6.16 ± 0.09 a | 1.91 ± 0.17 a | 0.65 ± 0.07 a | 0.47 ± 0.06 a | 0.24 ± 0.05 a |
| Medium-4 | 33.33 | 0.54 | 5.78 ± 0.21 a | 0.59 ± 0.07 a | 0 ± 0 a | 0 ± 0 a | 0 ± 0 a |
| Medium-5 | 4.44 | 1.08 | 2.54 ± 0.12 ab | 0 ± 0 a | 0 ± 0 a | 0 ± 0 a | 0 ± 0 a |
| Medium-6 | 11.11 | 1.08 | 3.57 ± 0.16 ab | 0 ± 0 a | 0 ± 0 a | 0 ± 0 a | 0 ± 0 a |
| Medium-7 | 17.77 | 1.08 | 2.10 ± 0.14 ab | 0 ± 0 a | 0 ± 0 a | 0 ± 0 a | 0 ± 0 a |
| Medium-8 | 33.33 | 1.08 | 1.43 ± 0.21 b | 0 ± 0 a | 0 ± 0 a | 0 ± 0 a | 0 ± 0 a |

Data represent mean ± standard errors (n = 10). Different letters followed by the means represent statistical differences at $p < 0.05$

## Discussion

It was Prof. E. C. Cocking whose pioneering works about plant protoplasts demonstrated this technique as a powerful tool for gene manipulation study in the plant. In 1960, Prof. Cocking published the first-ever report on plant protoplast isolation from the tomato root tip [32], which opened a new door to plant research. Soon after this, many plants were found feasible for protoplast study. There was a tremendous up-rise of protoplast research in the seventies and eighties. These days, protoplast systems are used as biotechnological tools for plant improvement by studying fundamental physiologic, metabolic, and developmental processes, transgenesis, gene manipulation, and somatic hybrid creation [2, 34, 35]. Despite having a superior oil quality, meagre oil yield is the primary concern that needs to be addressed for the successful commercialisation of “The Queen of Oilseed” sesame. Except for the three published reports, limited knowledge of sesame protoplast restricts the use of this potential tool for improving sesame. Shoji et al. [27], Bapat et al. [28], and Dhingra and Batra [29] reported the isolation and purification of sesame protoplast. Hypocotyls and calluses of sesame were the choice of explants in all these studies. Shoji et al. [27] and Bapat et al. [28] reported the protoplast division. For the first time, we are reporting a detailed step-wise protocol standardisation for protoplast isolation from hypocotyl, internode, leaf, and callus of sesame along with the micro-callus induction from hypocotyl-derived callus.

The enzymatic method of protoplast isolation is broadly dependent on three parameters – 1) *physical parameters:* explant preparation, and enzyme incubation period temperature, duration and shaking condition; 2) *chemical parameters:* selection and combination of enzymes, choice of osmoticum and its concentration, pretreatments, and additives; 3) *biological parameters:* plant species, explant type, and explant age. Protoplast isolation is a delicate task and a perfect balance between the three abovementioned parameter categories is an absolute mandate for success [35–38]. Some of these previously optimised protoplast isolation parameters of major oilseeds are enlisted in Table 5. In this study, we identified explant-wise differential optima of these parameters.

**Table 5.**
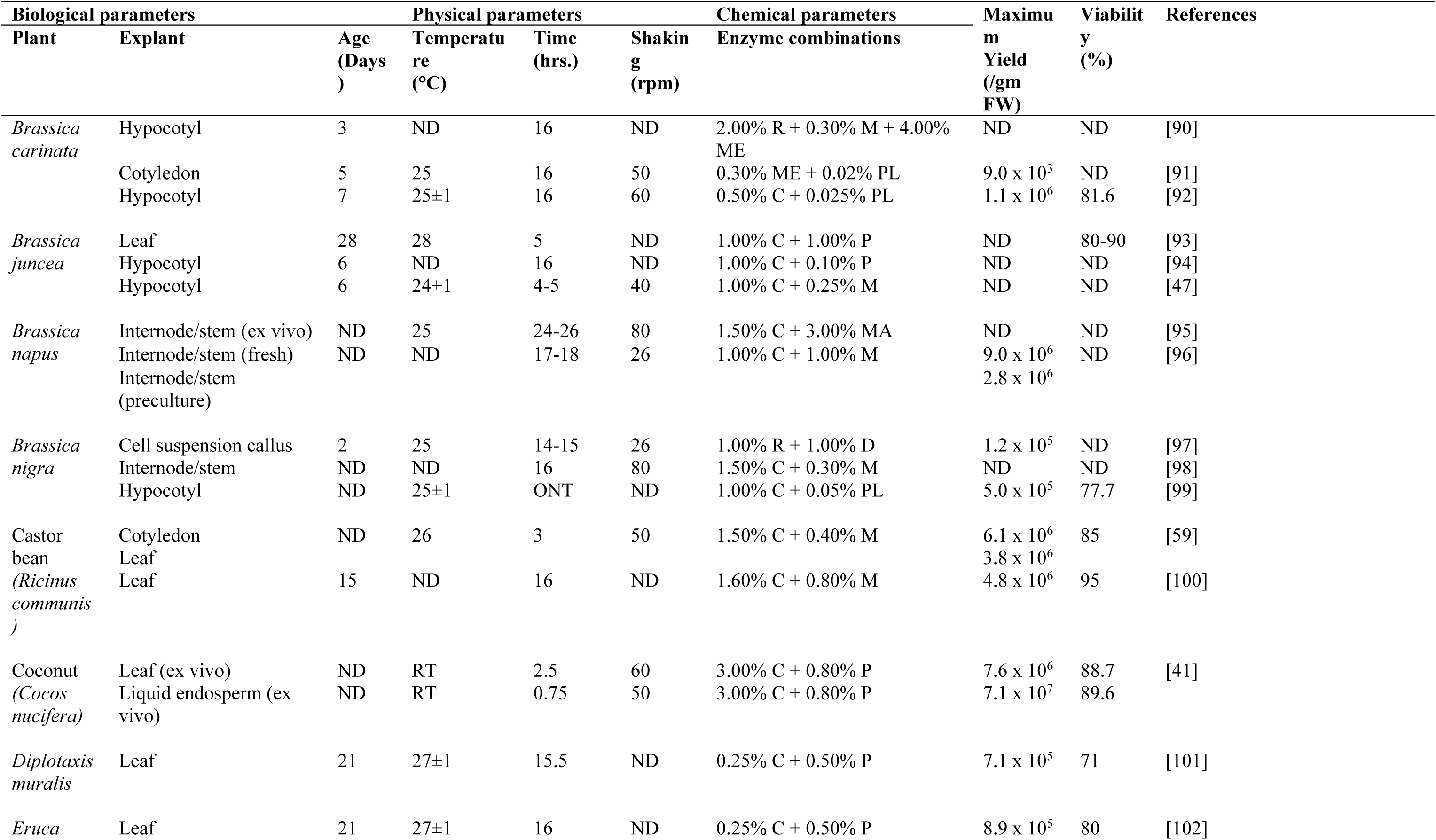
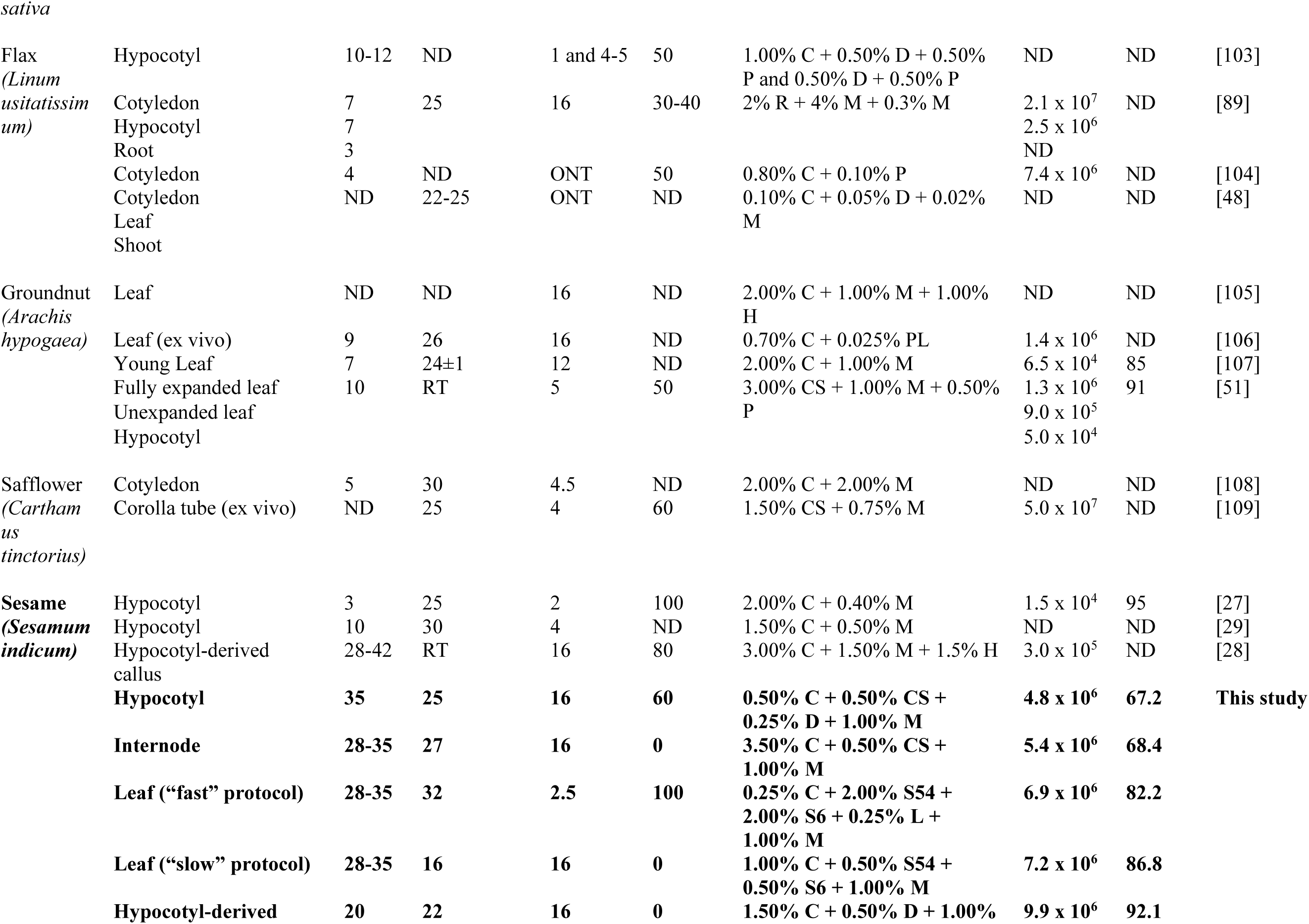
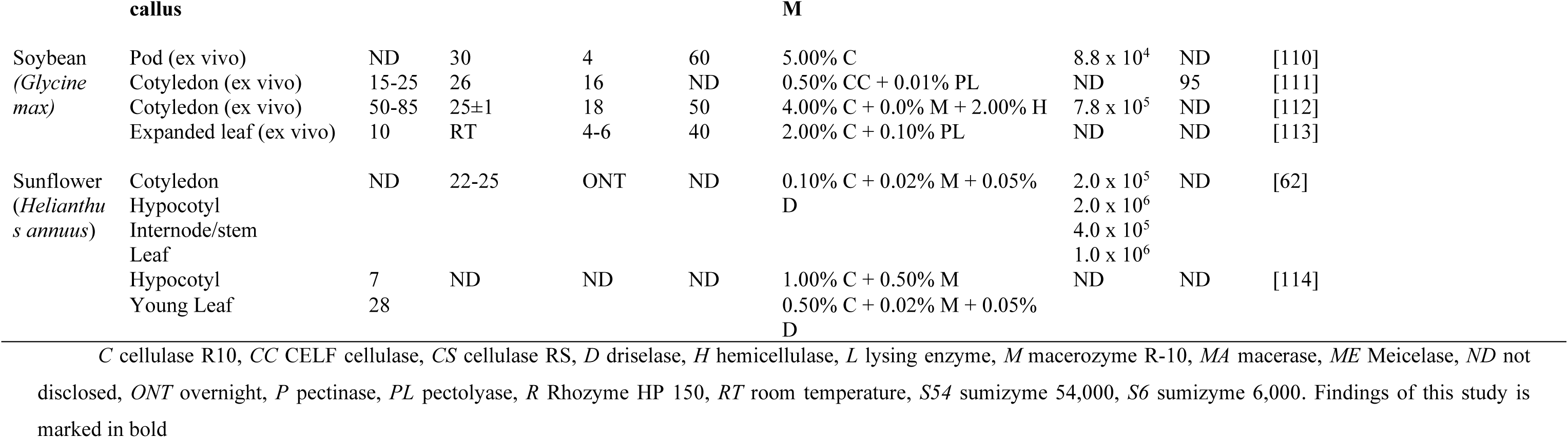
Different parameters of protoplast isolation from a few important oilseed plants.

### Optimisation of physical parameters for sesame protoplast isolation

Our study revealed an explant-wise combinational effect of enzyme incubation temperature and duration on protoplast yield and viability. We successfully isolated protoplasts from hypocotyl, internode, leaf (using “fast” protocol), leaf (using “slow” protocol), and callus explants using 25°C, 27°C, 32°C, 16°C, and 22°C temperatures, and 16 hrs, 16 hrs, 2.5 hrs, 16 hrs, and 16 hrs. duration for enzyme incubation, respectively (Table 1). These parameters vary widely among source plant species and tissue types. In *Medicago sativa*, enzyme incubation longer than six hrs. reduced protoplast yield and viability and increased the risk of enzyme toxicity-mediated protoplast death upon culturing; shorter incubation periods also diminished protoplast yield by incomplete plant tissue digestion [36]. Moon et al. [39] optimally isolated potato mesophyll protoplasts using 18 hrs. enzyme incubation. They also reported that five hrs. incubation decreased protoplast yields while 24 hrs. incubation decreased protoplast viability. Incubation at higher than optimal temperature affects enzyme activity and protoplast division. Shao et al. [37] demonstrated 26°C to be the optimum temperature to isolate protoplasts from *Uncaria rhynchophylla* leaves. Although the yield increased at higher than 26°C incubation temperatures, they obtained fewer active protoplasts. [37]. The combined effect of incubation temperature and duration on protoplast isolation was evident in many studies. Jeong et al. [19] described successful protoplast isolation from 10-day-old *Arabidopsis* seedlings using 12 hrs. enzyme incubation at 22-23°C. They obtained fragile protoplasts with low divisional response using >16 hrs. incubation. Protoplasts were optimally isolated from cultured young sporophytes of *Ecklonia cava* after 4 hrs. enzyme incubation at 20°C [40]. Xue et al. [38] reported that 16 hrs. enzymatic incubation at room temperature was optimum for *Duboisia* mesophyll protoplasts isolation. Previously, sesame protoplasts were isolated from hypocotyl explants using incubation at 25°C for 2 hrs. [27], and 30°C for 4 hrs. [29]. Bapat et al. [28] isolated suspension culture callus-derived protoplasts from seven sesame varieties at room temperature incubation for 16 hrs.

In our study, incubation shaking has an explant-wise differential effect on protoplast yield and viability. Shaking benefitted protoplast isolation from hypocotyls and leaves using the “fast” protocol. On the contrary, we found static incubation was best for protoplast isolation from internodes, leaves using the “slow” protocol, and calluses (Table 2). Due to the variability of cell wall composition among tissue types, the necessity and speed of incubational shaking depend on explants [35, 41]. Shaking during the enzymatic incubation period aids protoplast isolation by efficiently mixing enzyme solution with plant tissue [42]. However, due to the fragile nature of protoplasts, shaking speed needs to be optimised carefully to avoid any mechanical damage to the protoplasts. Generally, a maximum of 100 rpm shaking speed is applied for protoplast isolation (Table 5). High-speed shaking could disrupt chloroplast orientation and destroy protoplasts by cell membrane rupture [36]. Although some studies showed lower shaking speed or static incubation prolonged incubation duration and subsequent enzyme toxicity in protoplasts [19, 43], success reports of protoplasts isolation using static incubation do also exist [44–46, 38]. Previously for sesame protoplast isolation, Shoji et al. [27] and Bapat et al. [28] used shaking incubation, but Dhingra and Batra [29] did not use it.

### Optimisation of chemical parameters for sesame protoplast isolation

We established explant-wise differential optima of enzyme combinations in this study using commercially available cell wall degrading enzymes such as cellulase R-10, cellulase RS, driselase, lysing enzyme, macerozyme R-10, sumizyme 6,000, and sumizyme 54,000 (Table 3). Multiple studies have reported cell wall degradation by these enzymes [27–29, 47–49, 50–53]. Each enzyme possesses a variable amount of cellulolytic, hemicellulolytic, and pectinolytic activity. Plant cell walls are complex networks of cellulose, hemicelluloses, and pectins. Hydrolytic enzymes such as cellulases, hemicellulases, and pectinases can degrade respective cell wall components. A mixture of these enzymes could digest the cell wall completely [54]. Cellulose is a glucose homopolysaccharide. The three components of cellulase, endoglucanase, exoglucanase, and β-glucosidase synergistically hydrolyse cellulase into glucose [55]. Hemicelluloses are branched heteropolysaccharides of various hexoses and pentoses [54]. Hemicellulases act as glycoside hydrolases and carbohydrate esterases to break down hemicellulose into its monosaccharide components [56]. Pectins of plant cell walls are a diverse group of polysaccharides, consisting of galacturonic acid containing various functional groups and side chain mono- and polysaccharide components [54]. Pectinases degrade pectin through hydrolysis, trans-elimination, and de-esterification. During protoplast isolation, pectinases detach individual plant cells [57, 35]. Greatly variable cell wall composition among plant species and tissue types critically necessitates careful combination and concentration of these cell wall-degrading enzymes for successful protoplast isolation [35].

### Optimisation of biological parameters for sesame protoplast isolation

We have successfully isolated protoplasts from hypocotyls, internode, leaf, and callus explants. Protoplast yield and viability from calluses were highest among our tried explants (Table 3). Protoplast isolation from different explants of the same plant has been reported in apple [58], castor bean [59], coconut [41], *Cymbidium* sp. [60], flax [48, 89], groundnut [51], *Jasminum* sp. [61], sunflower [62], and tea [63]. Hypocotyl [27, 29], and callus [28] were the choice of explants for previous sesame protoplast studies. Our study is the first report of protoplast isolation from sesame internode and leaf explants. In the optimised protocol of this study, we achieved 9.88 ± 0.19 × 10^6^ protoplasts/gm FW yield and 92.05 ± 0.81% viability from callus explants. The yield is higher than the previous report [28]. However, the viability could not surpass the report of Shoji et al. [27] (95%).

Although it is a common belief that plant protoplasts behave indifferently regardless of the source tissue, Faraco et al. [64] argued on this point. Their study interestingly demonstrated source tissue-dependent differential tissue-specific promoter activity and subcellular protein localisation in petunia protoplasts. In another study, Di Sansebastiano et al. [65] used distinct membrane markers to identify different types of leaf cells. Therefore, our tissue-specific protoplast isolation protocol could benefit future spatial genomic and proteomic studies on sesame. We also believe that in future research, this protocol could be utilised as a single-cell system to investigate transient expression, subcellular localisation, protein-protein interaction, transcriptional regulation, CRISPR-based gene editing efficiency estimation, and many other omics and plant cellular process analyses.

### Regeneration of sesame callus-derived protoplasts to micro-callus

We observed a microtubule-like structure in cultured protoplasts before the 2-cell stage (Fig. 4a). This structure could be the mitotic prophase band indicating cell division initiation [66–68, 115]. Our callus-derived protoplasts divided and formed micro-callus (0.24 ± 0.05%) in BAP and NAA-fortified MS broth medium (Fig. 4; Table 4). Only one study demonstrated sesame protoplast-derived micro-callus formation to date, although the divisional rate was omitted in that report [28]. Combinations of BAP and NAA are used to regenerate callus-derived protoplasts in other plants; however, the divisional response of our sesame protoplasts was minuscule compared to these findings [69–72]. Neither the protoplasts derived from hypocotyl, internode, and leaf explants divided beyond the 2-cell stage nor the callus-derived protoplasts regenerated further than the micro-callus stage in our study. Such explant type dependency of protoplast regeneration has been described earlier [73–75]. Previous studies also depicted protoplast regeneration up to the micro-callus stage in *Brassica napus* [76], cabbage and cauliflower [77], *Cannabis sativa* [78], maize [79], and *Platycodon grandiflorus* [80].

A successful protoplast regeneration system demands critically optimised culture media and culture conditions [2]. The cell wall removal during protoplast isolation generates tremendous stress in plant cells [81–83]. Furthermore, commercial cell wall digesting enzymes act as stress elicitors generating ROS in isolated protoplasts [84]. Such a physiologic stressful condition often contributes to protoplast regeneration recalcitrancy. Fortification of culture media with antioxidants such as ascorbic acid, citric acid, reduced glutathione, and L-cysteine could mitigate stress and promote protoplast division [2, 74]. A well-balanced PGRs and osmoticum supplementations in culture media are also crucial for protoplast division and subsequent regeneration [2]. Various culture media additives such as activated charcoal, amino acids, arabinogalactan protein-rich extracts, complex organics (casein hydrolysate, casamino acids, coconut water, and yeast extract), galactoglucomannan-derived oligosaccharides, peptide growth factors (phytosulfokine), polyamines, polyvinylpyrrolidone, and silver nitrate are potential substances for protoplast regeneration success [82, 85–87, 2]. Since sesame in vitro regeneration is genotype-dependent, the choice of genotype might affect protoplast regeneration [88]. Finally, culturing conditions such as the plating density, the physical state of media (liquid or agar, agarose, alginate-based semi-solid), media refreshment interval, aeration, temperature, and the light-dark cycle can also contribute to this process [2, 74]. Therefore, a thorough investigation of these abovementioned factors could identify the reasons for low protoplast regeneration frequency and the caseation of micro-callus growth, and could potentially establish an efficient plant regeneration system from sesame protoplasts in future research.

## Conclusion

Herewith, we report an efficient protoplast isolation protocol utilising hypocotyl, internode, leaf, and callus explants of sesame. The maximum protoplast yield was obtained using callus explants and to date, it is the highest (9.88 ± 0.19 × 10^6^ protoplasts/gm FW) reported in sesame. Physical parameters such as enzyme incubation temperature, duration and shaking, chemical parameters such as enzyme combinations and concentrations, and biological parameters such as explant types played pivotal roles in protoplast yield and viability. We provided a reproducible, detailed, and stepwise protocol using a flow diagram from explants to protoplast plating. The micro-callus stage was obtained by callus-derived protoplast regeneration. The protoplast isolation system could be deployed to study various functional genomic, proteomic, cellular, and transient expression analyses leading to sesame development. Further investigation is required to achieve complete regenerated plants from sesame protoplast-derived micro-calluses.

## Abbreviations

2,4-D: 2,4-Dichlorophenoxyacetic acid
ANOVA: Analysis of Variance
BAP: 6-Benzylaminopurine
CRISPR: Clustered regularly interspaced short palindromic repeats
FDA: Fluorescein diacetate
FW: Fresh weight
MS: Murashige and Skoog
MES: 4-morpholineethanesulfonic acid
NAA: 1-Naphthaleneacetic acid
PGR: Plant growth regulator
RPM: Revolutions per minute
SE: Standard error

## Conflict of interest statement

The authors declare that they have no conflict of interest.

## Author contribution

Debabrata Basu and Samir Ranjan Sikdar arranged the funds and resources for the research, designed the experiments and supervised the work; Anirban Jyoti Debnath, Debabrata Basu, and Samir Ranjan Sikdar performed the experiments; Anirban Jyoti Debnath analysed the data, prepared the tables, graphics, and first draft of the manuscript. All the authors revised and approved the final manuscript.

## Statements and Declarations

The authors disclose no financial or non-financial interests directly or indirectly related to the work submitted for publication. They also have no relevant competing interests to declare to the content of this article.

## Funding

This work was supported by the National Agricultural Innovation Project, Indian Council of Agricultural Research (ICAR-NAIP) under Grant [Funding Number: NAIP/C4/C1090].

## Notes

### Competing Interest Statement

The authors have declared no competing interest.

### Summary of Updates

This version has revised author affiliation

## References

1. Avila-Peltroche J, Won BY, Cho TO (2022) An improved protocol for protoplast production, culture, and whole plant regeneration of the commercial brown seaweed *Undaria pinnatifida*. Algal Res 67:102851. 10.1016/j.algal.2022.102851

2. Reed KM, Bargmann BOR (2021) Protoplast Regeneration and Its Use in New Plant Breeding Technologies. Front Genome Ed 3:734951. 10.3389/fgeed.2021.734951

3. Nanjareddy K, Arthikala MK, Blanco L, Arellano ES, Lara M (2016) Protoplast isolation, transient transformation of leaf mesophyll protoplasts and improved *Agrobacterium*-mediated leaf disc infiltration of *Phaseolus vulgaris*: tools for rapid gene expression analysis. BMC Biotechnol 16(1):53. 10.1186/s12896-016-0283-8

4. Adjei MO, Zhao H, Tao X, Yang L, Deng S, Li X, Mao X, Li S, Huang J, Luo R, Gao A, Ma J (2023) Using A Protoplast Transformation System to Enable Functional Studies in *Mangifera indica* L. Int J Mol Sci 24(15):11984. 10.3390/ijms241511984

5. Kim K, Ho TS, Hwang US (2022) Production and characterization of somatic hybrids between rice (*Oryza sativa* L.) and reed (*Phragmites communis* Trin.) obtained by protoplast fusion. J Plant Biochem Biotechnol 31:370–379. 10.1007/s13562-021-00689-7

6. Ranaware AS, Kunchge NS, Lele SS, Ochatt SJ (2023) Protoplast Technology and Somatic Hybridisation in the Family Apiaceae. Plants (Basel) 12(5):1060. 10.3390/plants12051060

7. Goodman CD, Casati P, Walbot V (2004) A multidrug resistance-associated protein involved in anthocyanin transport in *Zea mays*. Plant Cell 16(7):1812–1826. 10.1105/tpc.022574

8. Lin HY, Chen JC, Fang SC (2018) A Protoplast Transient Expression System to Enable Molecular, Cellular, and Functional Studies in *Phalaenopsis* orchids. Front Plant Sci 9:843. 10.3389/fpls.2018.00843

9. Gilliard G, Huby E, Cordelier S, Ongena M, Dhondt-Cordelier S, Deleu M (2021) Protoplast: A Valuable Toolbox to Investigate Plant Stress Perception and Response. Front Plant Sci 12:749581. 10.3389/fpls.2021.749581

10. Sheen J (2001) Signal transduction in maize and Arabidopsis mesophyll protoplasts. Plant Physiol 127(4):1466–1475. 10.1104/pp.010820

11. Xing T, Wang X (2015) Protoplasts in plant signaling analysis: moving forward in the omics era. Botany 93(6): 325–332. 10.1139/cjb-2014-0219

12. Priyadarshani SVGN, Hu B, Li W, Ali H, Jia H, Zhao L, Ojolo SP, Azam SM, Xiong J, Yan M, Ur Rahman Z, Wu Q, Qin Y (2018) Simple protoplast isolation system for gene expression and protein interaction studies in pineapple (*Ananas comosus* L.). Plant Methods 14:95. 10.1186/s13007-018-0365-9

13. Liu Q, Li P, Cheng S, Zhao Z, Liu Y, Wei Y, Lu Q, Han J, Cai X, Zhou Z, Umer MJ, Peng R, Zhang B, Liu F (2022) Protoplast Dissociation and Transcriptome Analysis Provides Insights to Salt Stress Response in Cotton. Int J Mol Sci 23(5):2845. 10.3390/ijms23052845

14. Jiang F, Zhu J, Liu HL (2013) Protoplasts: a useful research system for plant cell biology, especially dedifferentiation. Protoplasma 250(6):1231–1238. 10.1007/s00709-013-0513-z

15. Xu Y, Li R, Luo H, Wang Z, Li MW, Lam HM, Huang C (2022) Protoplasts: small cells with big roles in plant biology. Trends Plant Sci 27(8):828–829. 10.1016/j.tplants.2022.03.010

16. Debnath AJ, Basu D, Sikdar SR (2014) An approach to standardise transient expression of GUS in sesame (*Sesamum indicum* L.) using Sonication Assisted *Agrobacterium tumefaciens*-mediated Transformation (SAAT) method. Isr J Plant Sci 61(1-4):37–45. 10.1080/07929978.2014.948275

17. Parveen S, Mazumder M, Bhattacharya A, Mukhopadhyay S, Saha U, Mukherjee A, Mondal B, Debnath AJ, Das S, Sikdar SR, Basu D. 2018. Identification of Anther-Specific Genes from Sesame and Functional Assessment of the Upstream Region of a Tapetum-Specific *β-1*, *3*-glucanase Gene. Plant Mol Biol Rep 36(2):149–161. 10.1007/s11105-017-1054-y

18. Nagata T, Takebe I (1971) Plating of isolated tobacco mesophyll protoplasts on agar medium. Planta 99(1):12–20. 10.1007/BF00392116

19. Jeong YY, Lee HY, Kim SW, Noh YS, Seo PJ (2021) Optimization of protoplast regeneration in the model plant Arabidopsis thaliana. Plant Methods 17(1):21. 10.1186/s13007-021-00720-x

20. Toriyama K, Hinata K (1985) Cell suspension and protoplast culture in rice. Plant Sci 41(3):179–183. 10.1016/0168-9452(85)90086-X

21. Webb KJ, Woodcock S, Chamberlain DA (1987) Plant Regeneration from Protoplasts of *Trifolium repens* and *Lotus corniculatus*. Plant Breed 98(2):111–118. 10.1111/j.1439-0523.1987.tb01102.x

22. Myers JR, Grosser JW, Taylor NL, Collins GB (1989) Genotype-dependent whole plant regeneration from protoplasts of red clover (*Trifolium pratense* L.). Plant Cell Tiss Organ Cult 19(2):113–127. 10.1007/BF00035811

23. Hayashimoto A, Li Z, Murai N (1990) A polyethylene glycol-mediated protoplast transformation system for production of fertile transgenic rice plants. Plant Physiol 93(3):857–63. 10.1104/pp.93.3.857

24. Marsan PA, Lupotto E, Locatelli F, Qiao Y, Cattaneo M (1993) Analysis of stable events of transformation in wheat via PEG-mediated DNA uptake into protoplasts. Plant Sci 93(1-2):85–94. 10.1016/0168-9452(93)90037-Z

25. Rhodes CA, Gray DW (1992) Transformation and regeneration of maize protoplasts. In: Lindsey K (ed) Plant Tissue Culture Manual. Springer, Boston, MA, pp 79–91. 10.1007/978-1-4899-3778-0_4

26. Monteiro M, Appezzato-da-Glória B, Valarini MJ, Alberto de Oliveira C, Vieira MLC (2003) Plant regeneration from proroplasts of alfalfa (*Medicago sativa*) via somatic embryogenesis. Sci agric (Piracicaba, Braz) 60(4):683–689. 10.1590/S0103-90162003000400012

27. Shoji K, Masuda K, Sugai M, Kobayashi T (1988) Relative DNA Content of Fused Protoplasts and Colony Cells in *Sesamum indicum*. Cytologia 53:205–211

28. Bapat VA, George L, Rao PS (1989) Isolation, culture and callus formation of sesame (*Sesamum indicum* L. cv. PT) protoplasts. Indian J Exp Biol 27:182–184

29. Dhingra M, Batra A (1990) High yielding preparation of viable protoplasts from hypocotyls of *Sesamum indicum* L. Curr Sci 59(6):325–326

30. Murashige T, Skoog S (1962) A revised medium for rapid growth and bioassay with tobacco tissue cultures. Physiol Plant 15:473–497. 10.1111/j.1399-3054.1962.tb08052.x

31. Power JB, Cocking EC (1970) Isolation of Leaf Protoplasts: Macromolecule Uptake and Growth Substance Response. J Exp Bot 21(1):64–70. 10.1093/jxb/21.1.64

32. Cocking EC (1960) A Method for the Isolation of Plant Protoplasts and Vacuoles. Nature 187:962–963. 10.1038/187962a0

33. Duncan DB (1955) Multiple range and multiple F test. Biometrics 11:1–42. 10.2307/3001478

34. Lin CS, Hsu CT, Yang LH, Lee LY, Fu JY, Cheng QW, Wu FH, Hsiao HC, Zhang Y, Zhang R, Chang WJ, Yu CT, Wang W, Liao LJ, Gelvin SB, Shih MC (2018a) Application of protoplast technology to CRISPR/Cas9 mutagenesis: from single-cell mutation detection to mutant plant regeneration. Plant Biotechnol J 16(7):1295–1310. 10.1111/pbi.12870

35. Chen K, Chen J, Pi X, Huang LJ, Li N (2023) Isolation, Purification, and Application of Protoplasts and Transient Expression Systems in Plants. Int J Mol Sci 24(23):16892. 10.3390/ijms242316892

36. Sangra A, Shahin L, Dhir S (2019) Optimization of Isolation and Culture of Protoplasts in Alfalfa (*Medicago sativa*) Cultivar Regen-SY. Am J Plant Sci 10(7):1206–1219. 10.4236/ajps.2019.107086

37. Shao Y, Mu D, Pan L, Wilson IW, Zheng Y, Zhu L, Lu Z, Wan L, Fu J, Wei S, Song L, Qiu D, Tang Q (2023) Optimization of Isolation and Transformation of Protoplasts from *Uncaria rhynchophylla* and Its Application to Transient Gene Expression Analysis. Int J Mol Sci 24(4):3633. 10.3390/ijms24043633

38. Xue Y, Hiti-Bandaralage JCA, Hu Z, Zhao Z, Mitter N (2024) First Report on Mesophyll Protoplast Isolation and Regeneration System for the *Duboisia* Species. Plants (Basel) 13(1):40. 10.3390/plants13010040

39. Moon KB, Park JS, Park SJ, Lee HJ, Cho HS, Min SR, Park YI, Jeon JH, Kim HS (2021) A More Accessible, Time-Saving, and Efficient Method for In Vitro Plant Regeneration from Potato Protoplasts. Plants (Basel) 10(4):781. 10.3390/plants10040781

40. Choi GC, Avila-Peltroche J, Won YB, Cho TO (2024) Optimization of protoplast isolation and subsequent regeneration from the economically important brown alga *Ecklonia cava* using response surface methodology. Algal Res 80:103525. 10.1016/j.algal.2024.103525

41. Guo Q, Wang Y, Zou J, Jing H, Li D (2023) Efficient isolation and transformation of protoplasts in coconut endosperm and leaves for gene function studies. Trop Plants 2:16 10.48130/TP-2023-0016

42. El-Gioushy SFEE, Kareem A, Baiea MHM (2019). Pre-isolation, isolation and regeneration protoplasts from leaf mesophyll of in vivo *Malus domestica* ‘Anna’ cv. Rev Bras Frutic 41(4):e-561. 10.1590/0100-29452019561

43. Cheng N, Nakata PA (2020) Development of a rapid and efficient protoplast isolation and transfection method for chickpea (*Cicer arietinum*). MethodsX 7:101025. 10.1016/j.mex.2020.101025

44. Blackhall NW, Davey MR, Power JB (1995) Isolation, culture, and regeneration of protoplasts. In: Dixon RA, Gonzales RA (eds) Plant Cell Culture: A Practical Approach. Oxford University Press, Oxford, UK

45. Babaoğlu M (2000) Protoplast Isolation in Lupin (*Lupinus mutabilis* Sweet): Determination of Optimum Explant Sources and Isolation Conditions. Turk J Bot 24(3):1777–185

46. Nassour M, Chassériaux G, Dorion N (2003) Optimization of protoplast-to-plant system for *Pelargonium*×*hortorum* ‘Alain’ and genetic stability of the regenerated plants. Plant Sci 165(1):121–128. 10.1016/S0168-9452(03)00150-X

47. Pua EC (1990) Somatic embryogenesis and plant regeneration from hypocotyl protoplasts of *Brassica juncea* (L.) Czern & Coss. Plant Sci. 68(2):231–238. 10.1016/0168-9452(90)90229-H

48. Bretagne-Sagnard B, Fouilloux G, Chupeau Y (1996) Induced albina mutations as a tool for genetic analysis and cell biology in flax *(Linum usitatissimum*), J Exp Bot 47(2):189–194. 10.1093/jxb/47.2.189

49. Rosnow J, Yerramsetty P, Berry JO, Okita TW, Edwards GE (2014) Exploring mechanisms linked to differentiation and function of dimorphic chloroplasts in the single cell C4 species *Bienertia sinuspersici*. BMC Plant Biol 14:34. 10.1186/1471-2229-14-34

50. Zaafouri K, Ziadi M, Ben Hassen-Trabelsi A, Mekni S, Aïssi B, Alaya M, Bergaoui L, Hamdi M (2017) Optimization of Hydrothermal and Diluted Acid Pretreatments of Tunisian *Luffa cylindrica* (L.) Fibers for 2G Bioethanol Production through the Cubic Central Composite Experimental Design CCD: Response Surface Methodology. Biomed Res Int 2017:9524521. 10.1155/2017/9524521

51. Biswas S, Wahl NJ, Thomson MJ, Cason JM, McCutchen BF, Septiningsih EM (2022) Optimization of Protoplast Isolation and Transformation for a Pilot Study of Genome Editing in Peanut by Targeting the Allergen Gene *Ara h 2*. Int J Mol Sci 23(2):837. 10.3390/ijms23020837

52. Ning Y, Hu B, Yu H, Liu X, Jiao B, Lu X (2022) Optimization of Protoplast Preparation and Establishment of Genetic Transformation System of an Arctic-Derived Fungus *Eutypella* sp. Front Microbiol 13:769008. 10.3389/fmicb.2022.769008

53. Yang P, Sun Y, Sun X, Li Y, Wang L (2024) Optimization of preparation and transformation of protoplasts from *Populus simonii* × *P. nigra* leaves and subcellular localization of the major latex protein 328 (MLP328). Plant Methods 20(1):3. 10.1186/s13007-023-01128-5

54. de Souza TSP, Kawaguti HY (2021) Cellulases, Hemicellulases, and Pectinases: Applications in the Food and Beverage Industry. Food Bioprocess Technol 14:1446–1477. 10.1007/s11947-021-02678-z

55. Ejaz U, Sohail M, Ghanemi A (2021) Cellulases: From Bioactivity to a Variety of Industrial Applications. Biomimetics (Basel) 6(3):44. 10.3390/biomimetics6030044

56. Shallom D, Shoham Y (2003) Microbial hemicellulases. Curr Opin Microbiol 6(3):219–228. 10.1016/S1369-5274(03)00056-0

57. Sakai T, Sakamoto T, Hallaert J, Vandamme EJ (1993) Pectin, pectinase and protopectinase: production, properties, and applications. Adv Appl Microbiol 39:213–294. 10.1016/s0065-2164(08)70597-5

58. Ai-ping D, Hong-fan E, Yu-fen C (1994) Protoplast Culture and Plant Regeneration of *Malus pumila*. Acta Bot Sin 36(4):271–277

59. Bai L, Cheng Y, She J, He Z, Liu H, Zhang G, Cao R (2020) Development of an efficient protoplast isolation and transfection system for castor bean (*Ricinus communis* L.). Plant Cell Tiss Organ Cult 143:457–464. 10.1007/s11240-020-01932-0

60. Ren R, Gao J, Yin D, Li K, Lu C, Ahmad S, Wei Y, Jin J, Zhu G, Yang F (2021) Highly Efficient Leaf Base Protoplast Isolation and Transient Expression Systems for Orchids and Other Important Monocot Crops. Front Plant Sci 12:626015. 10.3389/fpls.2021.626015

61. Ahmed MAA, Miao M, Pratsinakis ED, Zhang H, Wang W, Yuan Y, Lyu M, Iftikhar J, Yousef AF, Madesis P, Wu B (2021) Protoplast Isolation, Fusion, Culture and Transformation in the Woody Plant *Jasminum* spp. Agric 11(8):699. 10.3390/agriculture11080699

62. Lenee P, Chupeau Y (1986) Isolation and culture of sunflower protoplasts (*Helianthus annuus* L.): Factors influencing the viability of cell colonies derived from protoplasts. Plant Sci 43(1):69–75. 10.1016/0168-9452(86)90110-X

63. Xu XF, Zhu HY, Ren YF, Feng C, Ye ZH, Cai HM, Wan XC, Peng CY (2021) Efficient isolation and purification of tissue-specific protoplasts from tea plants (*Camellia sinensis* (L.) O. Kuntze). Plant Methods 17(1):84. 10.1186/s13007-021-00783-w

64. Faraco M, Di Sansebastiano GP, Spelt K, Koes RE, Quattrocchio FM (2011) One protoplast is not the other! Plant Physiol 156(2):474–478. 10.1104/pp.111.173708

65. Di Sansebastiano GP, Paris N, Marc-Martin S, Neuhaus JM (2001) Regeneration of a lytic central vacuole and of neutral peripheral vacuoles can be visualized by green fluorescent proteins targeted to either type of vacuoles. Plant Physiol 126(1):78–86. 10.1104/pp.126.1.78

66. Simmonds DH (1986) Prophase bands of microtubules occur in protoplast cultures of *Vicia hajastana* Grossh. Planta. 167(4):469–472. 10.1007/BF00391222

67. Meijer EGM, Simmonds DH (1988) Microtubule organization during the development of the mitotic apparatus in cultured mesophyll protoplasts of higher plants – an immunofluorescence microscopic study. Physiol Plant 74(2):225–232. 10.1111/j.1399-3054.1988.tb00625.x

68. Meijer EGM, Simmonds DH (1989) Organization of Actin Microfilaments in Cultured Mesophyll Protoplasts of *Medicago sativa* and *Nicotiana tabacum*. J Plant Physiol 135(1):122–125. 10.1016/S0176-1617(89)80236-6

69. Dunbar KB, Stephens CT (1991) Plant regeneration from callus-derived protoplasts of *Pelargonium* x *domesticum*. Plant Cell Rep 10:417–420. 10.1007/BF00232615

70. Santos C, Caldeira G (1998) Callus formation and plant regeneration from protoplasts of sunflower calli and hypocotyls. Acta Societatis Botanicorum Poloniae 67(1):31–36. 10.5586/asbp.1998.003

71. Kanwar K, Bhardwaj A, Deepika R (2009) Efficient regeneration of plantlets from callus and mesophyll derived protoplasts of *Robinia pseudoacacia* L. Plant Cell Tiss Organ Cult 96:95–103. 10.1007/s11240-008-9465-y

72. Klimek-Chodacka M, Kadluczka D, Lukasiewicz A, Malec-Pala A, Baranski R, Grzebelus E (2020) Effective callus induction and plant regeneration in callus and protoplast cultures of *Nigella damascena* L. Plant Cell Tiss Organ Cult 143:693–707. 10.1007/s11240-020-01953-9

73. Roest S, Gilissen LJW (1989) Plant regeneration from protoplasts: a literature review. Acta Bot Neerl 38(1):1–23. 10.1111/j.1438-8677.1989.tb01907.x

74. Eeckhaut T, Lakshmanan PS, Deryckere D, Bockstaele EV, Huylenbroeck JV (2013) Progress in plant protoplast research. Planta 238:991–1003. 10.1007/s00425-013-1936-7

75. Rahmani MS, Pijut PM, Shabanian N (2016) Protoplast isolation and genetically true-to-type plant regeneration from leaf- and callus-derived protoplasts of *Albizia julibrissin*. Plant Cell Tiss Organ Cult 127:475–488. 10.1007/s11240-016-1072-8

76. Sun M, Kieft H, Zhou C, nvan Lammeren A (1999) A co-culture system leads to the formation of microcalli derived from microspore protoplasts of *Brassica napus* L. cv. Topas. Protoplasma 208:265–274. 10.1007/BF01279098

77. Hussain M, Li H, Badri Anarjan M, Lee S (2024) Development of a general protoplast-mediated regeneration protocol for *Brassica*: cabbage and cauliflower as examples. Hortic Environ Biotechnol 65:313–321. 10.1007/s13580-023-00557-4

78. Monthony AS, Jones AMP (2024) Enhancing Protoplast Isolation and Early Cell Division from *Cannabis sativa* Callus Cultures via Phenylpropanoid Inhibition. Plants 13(1):130. 10.3390/plants13010130

79. Imbrie-Milligan CW, Hodges TK (1986) Microcallus formation from maize protoplasts prepared from embryogenic callus. Planta 168:395–401. 10.1007/BF00392367

80. Kwon SH, Murthy HN, Han JE, Lee HS, Park SY (2024) Isolation, culture of *Platycodon grandiflorus* protoplasts: factors affecting protoplast yield, cell division, and micro-callus formation. Hortic Environ Biotechnol 65:515–525. 10.1007/s13580-023-00585-0

81. Cassells A, Curry R (2001) Oxidative stress and physiological, epigenetic and genetic variability in plant tissue culture: implications for micropropagators and genetic engineers. Plant Cell Tiss Org 64:145–157. 10.1023/A:1010692104861

82. Papadakis AK, Roubelakis-Angelakis KA (1999) The generation of active oxygen species differs in tobacco and grapevine mesophyll protoplasts. Plant Physiol 121(1):197–206. 10.1104/pp.121.1.197

83. Reyna-Llorens I, Ferro-Costa M, Burgess SJ (2023) Plant protoplasts in the age of synthetic biology. J Exp Bot 74(13):3821–3832. 10.1093/jxb/erad172

84. Ishii S (1988) Factors Influencing Protoplast Viability of Suspension-Cultured Rice Cells during Isolation Process. Plant physiol 88(1):26–29. 10.1104/pp.88.1.26

85. Wiśniewska E, Majewska-Sawka A (2007) Arabinogalactan-proteins stimulate the organogenesis of guard cell protoplasts-derived callus in sugar beet. Plant Cell Rep 26(9):1457–1467. 10.1007/s00299-007-0348-1

86. Kakoniova D, Hlinkova E, Liskova D, Kollarova K (2010) Oligosaccharides induce changes in protein patterns of regenerating spruce protoplasts. Cent Eur J Biol 5(3):353–363. 10.2478/s11535-010-0018-0

87. Grzebelus E, Szklarczyk M, Gren J, Sniegowska K, Jopek M, Kacinska I, Mrozek K (2012) Phytosulfokine stimulates cell divisions in sugar beet (*Beta vulgaris* L.) mesophyll protoplast cultures. Plant Growth Regul 67(1):93–100. 10.1007/s10725-011-9654-2

88. Debnath AJ, Gangopadhyay G, Basu D, Sikdar SR (2018) An efficient protocol for in vitro direct shoot organogenesis of *Sesamum indicum* L. using cotyledon as explant. 3 Biotech 8(3):146. 10.1007/s13205-018-1173-7

89. Barakat MN, Cocking EC (1983) Plant regeneration from protoplast-derived tissues of *Linum usitatissimum* L. (Flax). Plant Cell Rep 2(6):314–317. 10.1007/BF00270190

90. Chuong PV, Pauls KP, Beversdorf WD (1987) Protoplast culture and plant regeneration from *Brassica carinata* Braun. Plant Cell Rep 6:67–69. 10.1007/BF00269742

91. Jaiswal SK, Hammatt N, Bhojwani SS, Cocking EC, Davey MR (1990) Plant regeneration from cotyledon protoplasts of *Brassica carinata*. Plant Cell Tiss Organ Cult 22:159–165. 10.1007/BF00033630

92. Narasimhulu SB, Kirti PB, Prakash S, Chopra VL (1992) Rapid and efficient plant regeneration from hypocotyl protoplasts of *Brassica carinata*. Plant Cell Rep 11, 159–162 (1992). 10.1007/BF00232171

93. Chatterjee G, Sikdar SR, Das S, Sen SK (1985) Regeneration of plantlets from mesophyll protoplasts of *Brassica juncea* (L.) Czern. Plant Cell Rep 4(5):245–247. 10.1007/BF00269368

94. Kirti PB, Chopra VL (1989) Plant regeneration from hypocotyl-derived protoplasts of *Brassica juncea* (L.) Czern and Coss. Plant Cell Rep 7:708–710. 10.1007/BF00272067

95. Chuong PV, Beversdorf WD Pauls KP (1987) Plant Regeneration from Haploid Stem Peel Protoplasts of *Brassica napus* L. J Plant Physiol 130(1):57–65. 10.1016/S0176-1617(87)80301-2

96. Klimaszewska K, Keller WA (1987) Plant regeneration from stem cortex protoplasts of *Brassica napus*. Plant Cell Tiss Organ Cult 8:225–233. 10.1007/BF00040949

97. Klimaszewska K, Keller WA (1985) Somatic Embryogenesis in Cell Suspension and Protoplast Cultures of *Brassica nigra* (L.) Koch. J Plant Physiol 122(3):251–260. 10.1016/S0176-1617(86)80124-9

98. Chuong PV, Pauls KP, Beversdorf WD (1987) Plant regeneration from *Brassica nigra* (L.) Koch stem protoplasts. In Vitro Cell Dev Biol 23(6):449–452. 10.1007/BF02623862

99. Narasimhulu SB, Kirti PB, Prakash S, Chopra VL (1993) Rapid and high frequency shoot regeneration from hypocotyl protoplasts of *Brassica nigra*. Plant Cell Tiss Organ Cult 32:35–39. 10.1007/BF00040113

100. Figueroa-Varela P, Susunaga-Gómez D, Restrepo-Osorio C, Harms C, Villanueva-Mejía D (2023) An efficient method for protoplast-mediated production of transformed castor bean (*Ricinus communis*) lines. BMC Res Notes 16(1):140. 10.1186/s13104-023-06414-y

101. Sikdar SR, Sengupta S, Das S, Sen SK (1990). Plant regeneration from mesophyll protoplasts of *Diplotaxis muralis*, a wild crucifer. Plant Cell Rep 8(12):722–725. 10.1007/BF00272103

102. Sikdar SR, Chatterjee G, Das S, Sen SK (1987) Regeneration of plants from mesophyll protoplasts of the wild crucifer *Eruca sativa* Lam. Plant Cell Rep 6(6):486–489. 10.1007/BF00272790

103. Gamborg OL, Shyluk JP (1976) Tissue Culture, Protoplasts, and Morphogenesis in Flax. Bot Gaz 137(4): 301–306. 10.1086/336875

104. David H, David A, Bade P, Millet J, Morvan O, Morvan C (1994) Cell Wall Composition and Morphogenic Response in Callus Derived from Protoplasts of two Fibre flax (*Linum usitatissimum* L.) Genotypes. J Plant Physiol 43(3):379–384. 10.1016/S0176-1617(11)81648-2

105. Oelck MM, Bapat VA, Schieder O (1982) Protoplast Culture of Three Legumes: *Arachis hypogaea*, *Melilotus officinalis*, *Trifolium resupinatum*. Z Pflanzenphysiologie 106(2):173–177. 10.1016/S0044-328X(82)80080-9

106. Rugrnan EE, Cocking EC (1985) The development of somatic hybridization technique for groundnut improvement. In: Moss JP (ed) Proceedings of International Workshop on Cytogenetics of Arachis. 31 Oct-2 Nov 1983, ICRISAT, Patancheru, India, pp 167–174

107. Venkatachalam P, Jayabalan N (1996) Efficient callus induction and plant regeneration via somatic embryogenesis from immature leaf-derived protoplasts of groundnut (*Arachis hypogaea* L.). Isr J Plant Sci 44(4):387–396. 10.1080/07929978.1996.10676660

108. Ichihara K, Noda M (1981) Lipid synthesis in germinating safflower seeds and protoplasts. Phytochemistry 20(5):1023–1030. 10.1016/0031-9422(81)83021-X

109. Zhou YX, Wang J, Peng YN, Chen C, Xian B, Xi ZQ, Ren CX, Pei J, Chen J (2024) CtMYB1 regulate flavonoid biosynthesis in safflower flower by binding the CAACCA elements. Res Sq (preprint V1) 10.21203/rs.3.rs-4188109/v1

110. Zieg RG, Outka DE (1980) The isolation, culture and callus formation of soybean pod protoplasts. Plant Sci Lett 18(2):105–114. 10.1016/0304-4211(80)90038-3

111. Lin W, Odell JT, Schreiner RM (1987) Soybean Protoplast Culture and Direct Gene Uptake and Expression by Cultured Soybean Protoplasts. Plant Physiol 84(3):856–861. 10.1104/pp.84.3.856

112. Wei ZM, Xu ZH (1988) Plant regeneration from protoplasts of soybean (*Glycine max* L.). Plant Cell Rep 7:348–351. 10.1007/BF00269935

113. Wu F, Hanzawa Y (2018) A Simple Method for Isolation of Soybean Protoplasts and Application to Transient Gene Expression Analyses. J Vis Exp 131:57258. 10.3791/57258

114. Kativat C, Chueakhunthod W, Tantasawat PA (2022) The effects of cytokinin and plating density on protoplast culture of sunflower. J Plant Biotechnol 49(4):331–338. 10.5010/JPB.2022.49.4.331

115. Dijak M, Simmonds DH (1988) Microtubule organization during early direct embryogenesis from mesophyll protoplasts of *Medicago sativa* L. Plant Sci 58(2):183–191. 10.1016/0168-9452(88)90008-8

116. Widholm JM (1972) The use of fluorescein diacetate and phenosafranine for determining viability of cultured plant cells. Stain Technol 47(4):189–194. 10.3109/10520297209116483

