## Supplementary Table S1 for "Optimisation of explant-specific isolation, culture, and micro-callus induction of sesame (*Sesamum indicum* L.) protoplasts"

### ARTICLE TYPE

Full-Length Research Article

### AUTHOR INFORMATION

Anirban Jyoti Debnath[1]*, Debabrata Basu[1], and Samir Ranjan Sikdar[1]

[1] Division of Plant Biology, Bose Institute, Centenary Campus, P-1/12, C. I. T. Road, Scheme VII M, Kolkata – 700 054, West Bengal, India

*Corresponding Author ORCID ID: 0000-0003-1907-837X

**Supplementary Table S1** Enzyme combinations used to isolate protoplasts from different sesame explants in the first experiment

| Explants | Enzyme combinations |
|---|---|
| Hypocotyl | Cellulase R-10 0.5%<br>Cellulase RS 0.25%<br>Driselase 0.25%<br>Macerozyme R-10 1.0% |
| Internode | Cellulase R-10 0.5%<br>Cellulase RS 0.25%<br>Driselase 0.25%<br>Macerozyme R-10 1.0% |
| Leaf "fast" protocol | Cellulase R-10 1.0%<br>Sumizyme 54,000 1.0%<br>Sumizyme 6,000 0.5%<br>Lysing enzyme 0.25%<br>Macerozyme R-10 1.0% |
| Leaf "slow" protocol | Cellulase R-10 1.0%<br>Sumizyme 54,000 1.0%<br>Sumizyme 6,000 1.0%<br>Lysing enztme 1.0%<br>Macerozyme R-10 1.0% |
| Callus | Cellulase R-10 0.5%<br>Driselase 0.5%<br>Macerozyme R-10 1.0% |
